# Dynamin2 stabilizes plasma membrane-connected caveolae by restraining fission

**DOI:** 10.1101/2022.05.25.493372

**Authors:** Elin Larsson, Björn Morén, Kerrie-Ann McMahon, Robert G. Parton, Richard Lundmark

**Author notes:** Corresponding author: Richard Lundmark, Department of Integrative Medical Biology, Umeå University, 901 87 Umeå, Sweden.

## Abstract

Caveolae are small membrane invaginations that generally are stably attached to the plasma membrane. Their release is believed to depend on the GTPase dynamin 2 (Dyn2), in analogy with its role in fission of clathrin-coated vesicles. The mechanistic understanding of caveola fission, and in particular the role of Dyn2, is however sparse. Here, we used microscopy-based tracking of individual caveolae in living cells to determine the role of Dyn2 in caveola dynamics. We report that Dyn2 stably associated with a subset of caveolae, but was not required for formation or fission of caveolae. The Dyn2-positive subset of caveolae displayed longer plasma membrane duration times, whereas depletion of Dyn2 resulted in shorter duration times and increased caveola fission. The stabilizing role of Dyn2 was independent of its GTPase activity and the caveola stabilizing protein EHD2. Thus, we propose that, in contrast to the current view, Dyn2 restrains caveolae at the plasma membrane and prevents fission.

## Introduction

Membrane fission is essential for the release of membrane transport vesicles in the cell. Classically, vesicles associated with the membrane via a membrane pore or neck, has been seen as transient states in the process of vesicle release or fusion. However, several important processes rely on stabilization of the attached vesicle for an extended period of time (Neufeldt et al., 2018; Parton, 2018). Currently, it is largely unknown how this stabilisation and temporal restrain of fission is controlled in living cells. Caveolae are small bulb-shaped invaginations of the plasma membrane (PM) with atypical lipid composition and dynamics as compared to classical membrane vesicles such as clathrin- or COP-coated vesicles (Parton, 2018). Caveolae are characteristically constrained to the cell surface as fully invaginated buds, but can also flatten out (Sinha et al., 2011) or undergo short-range cycles of fission and fusion (Pelkmans and Zerial, 2005).

In non-muscle cells, caveola formation is dependent on lipid-driven assembly of the integral membrane protein caveolin1 (Cav1) (Monier et al., 1995; Murata et al., 1995; Rothberg et al., 1992) and the peripherally membrane attached protein cavin1(Aboulaich et al., 2004). Cav1 form complexes with cholesterol in the membrane to which cavin1 is recruited leading to invagination of such cholesterol-rich domains (Hill et al., 2008). The large ATPase EHD2, belonging to the dynamin superfamily of proteins, is present on most caveolae (Hansen et al., 2011; Moren et al., 2012; Stoeber et al., 2012). EHD2 self-oligomerizes at the caveolae neck, thereby preventing fission and constraining caveolae to the PM. Caveola stabilization is dependent on ATP binding and hydrolysis by EHD2 resulting in conformational changes which facilitate oligomerization into rings surrounding membranes tubes (Hoernke et al., 2017). Other proteins have also been shown to influence caveola dynamics. The BAR domain containing protein pacsin2 (Pac2) binds to EHD2 and appear to aid in the stabilization of caveolae (Moren et al., 2012; Senju et al., 2015). Additionally, caveolae are tightly coupled to actin filaments which could restrict both lateral movement and internalization (Echarri and Del Pozo, 2015). Yet, it is clear that caveolae also detach from the PM in most cell types. Different models for caveolae fission have been proposed including both lipid- and protein-driven mechanisms. Accumulation of cholesterol and glycosphingolipids were shown to drive caveola fission in a process which could be counteracted by EHD2 (Hubert et al., 2020). In addition, incorporation of cavin3 in the caveolae coat was shown to increase fission (Mohan et al., 2015), and coupling of caveolae to actin via filamin was shown to promote internalization (Stahlhut and van Deurs, 2000; Sverdlov et al., 2009). In analogy with its role in clathrin mediated endocytosis, dynamin 2 (Dyn2) has been proposed to perform fission of caveolae (Henley et al., 1998; Oh et al., 1998; Schnitzer et al., 1996).

Dyn2 is ubiquitously expressed and belongs to the dynamin superfamily of large GTPases involved in membrane fission and fusion processes. The homologues, dynamin 1 and 3 are primarily expressed in the brain. Dynamins are comprised of a G-domain that binds and hydrolyses GTP, the bundle signaling element (BSE), a helical stalk domain, a PH domain and a proline rich region that binds to SH3-domain containing proteins. Dynamins are known to form dimers and tetramers and to self-assemble into higher order structures such as rings and helices. The formation of helices around membrane tubes has been extensively studied in relation to the typical role of dynamins in catalyzing fission of clathrin coated vesicles (CCV) for endocytosis. In the GTP-bound state, the stalk and PH domain bind to membranes, which facilitate self-oligomerization of dynamins into short helical rings. Nucleotide-dependent conformational changes powers constriction of the helical polymer and the underlying membrane resulting in fission. GTP hydrolysis also mediates disassembly of the helices and membrane release of dynamins (Antonny et al., 2016).

It has become apparent that dynamins can also function via other mechanisms. For example, dynamin was also identified to play a key role in the early stages of CCV formation (Aguet et al., 2013; Reis et al., 2015) where oligomerization and assembly-stimulated GTPase activity are not required (Sever et al., 2000; Sever et al., 1999). Furthermore, dynamin influences a wide range of actin-driven processes (Sever et al., 2013), and binds directly to actin (Gu et al., 2010; Palmer et al., 2015) in a mechanism independent of oligomerization and GTP binding and hydrolysis. Furthermore, the dynamin scaffolds used for membrane constriction during fission could also stabilize membrane tubules and prevent fission (Bashkirov et al., 2008; Boucrot et al., 2012; Pucadyil and Schmid, 2008). Indeed, while dynamin 1 can perform scission of vesicles from planar lipid membranes, Dyn2 acts in synergy with other proteins to perform fission (Liu et al., 2011; Neumann and Schmid, 2013). In spite of the vast literature on the role of dynamin in membrane fission processes, little is known about its role at caveolae. Dyn2 is the dynamin homologue predominantly expressed in cells with caveolae. Cell free assays of caveolae budding based on purified membranes and cytosol suggested that GTP hydrolysis and dynamin was required for caveola fission (Oh et al., 1998; Schnitzer et al., 1996). Inhibition of cholera toxin B-subunit (CTxB) uptake by injection of antibodies against Dyn2 or expression of the GTPase deficient mutant K44A has been interpreted as an effect on caveolae fission (Henley et al., 1998). However, CTxB is mainly internalized by other pathways (Kalia et al., 2006; Kirkham et al., 2005; Lundmark et al., 2008; Massol et al., 2004; Torgersen et al., 2001) and the regulatory role of Cav1 in CTxB-uptake is independent of caveolae (Lajoie et al., 2009). Thus, many questions remain regarding the role of Dyn2 at caveolae and the fission of these atypical membrane invaginations.

Here, we have addressed the role of Dyn2 in caveola fission in living cells. Using a cell system with inducible expression of fluorescently labelled caveolae proteins we show that Dyn2 is stably associated with a distinct subset of caveolae. By Total Internal Reflection Fluorescence (TIRF) microscopy and particle tracking we have determined how caveola dynamics is influenced by Dyn2. We show that Dyn2 restrains caveolae fission and increase the time that they are associated with the plasma membrane. Increased levels of Dyn2 counteracts lipid-induced fission and this stabilizing effect is independent of its GTPase activity. Dyn2 acts independently of the caveola stabilizing protein EHD2 providing an additive mechanism to confine caveolae to the plasma membrane.

## Results

### Dyn2 depletion makes caveolae more dynamic in HeLa cells

In agreement with the proposed role of Dyn2 at caveolae (Oh et al., 1998), immunogold labeling of PM lawns of differentiated 3T3-L1 cells confirmed the presence of Dyn2 at a subset of the numerous caveolae in these cells (Fig. 1A). However, compared to the localization at clathrin coated pits, the Dyn2 labelling was sparse. In order to further investigate the proposed role of Dyn2 in caveolae fission, we tracked real time caveola dynamics using a HeLa Cav1-mCh FlpIn cell line (Hubert et al., 2020). In these cells, expression of mCherry-tagged caveolin1 (Cav1-mCh) is induced to near endogenous levels following addition of doxycycline. TIRF imaging allowed restricted visualization of caveolae at or in close proximity to the PM (Fig. 1B). TIRF movies were analyzed using the Imaris tracking software, were Cav1-positive spots of a certain size and fluorescent intensity were regarded as a caveolae (Fig. 1B and video 1). These caveolae were tracked over time and the dynamic parameters i) duration time, ii) displacement length and iii) mean speed was examined (Fig. 1B). By combining these parameters, we could follow a large number of caveolae and discriminate between stable and dynamic caveolae. Long duration times, short displacement length and low track mean speed are indicative of caveolae stably connected to the cell surface (Fig. 1B, line 1 and red symbols). Intermediate duration time, high speed and long displacement length is interpreted as fissioned caveolae moving rapidly in the plane of the membrane (Fig. 1B, line 2 and cyan symbols). Short duration times, short displacement length and high mean speed is indicative of fissioned caveolae moving rapidly in and out of the TIRF plane (Fig. 1B, line3 and 4 and purple and blue symbols). Tracking of the caveolae showed a heterogenous stability within a cell as has previously been described (Boucrot et al., 2011; Hoernke et al., 2017; Hubert et al., 2020; Mohan et al., 2015).

**Fig. 1.**
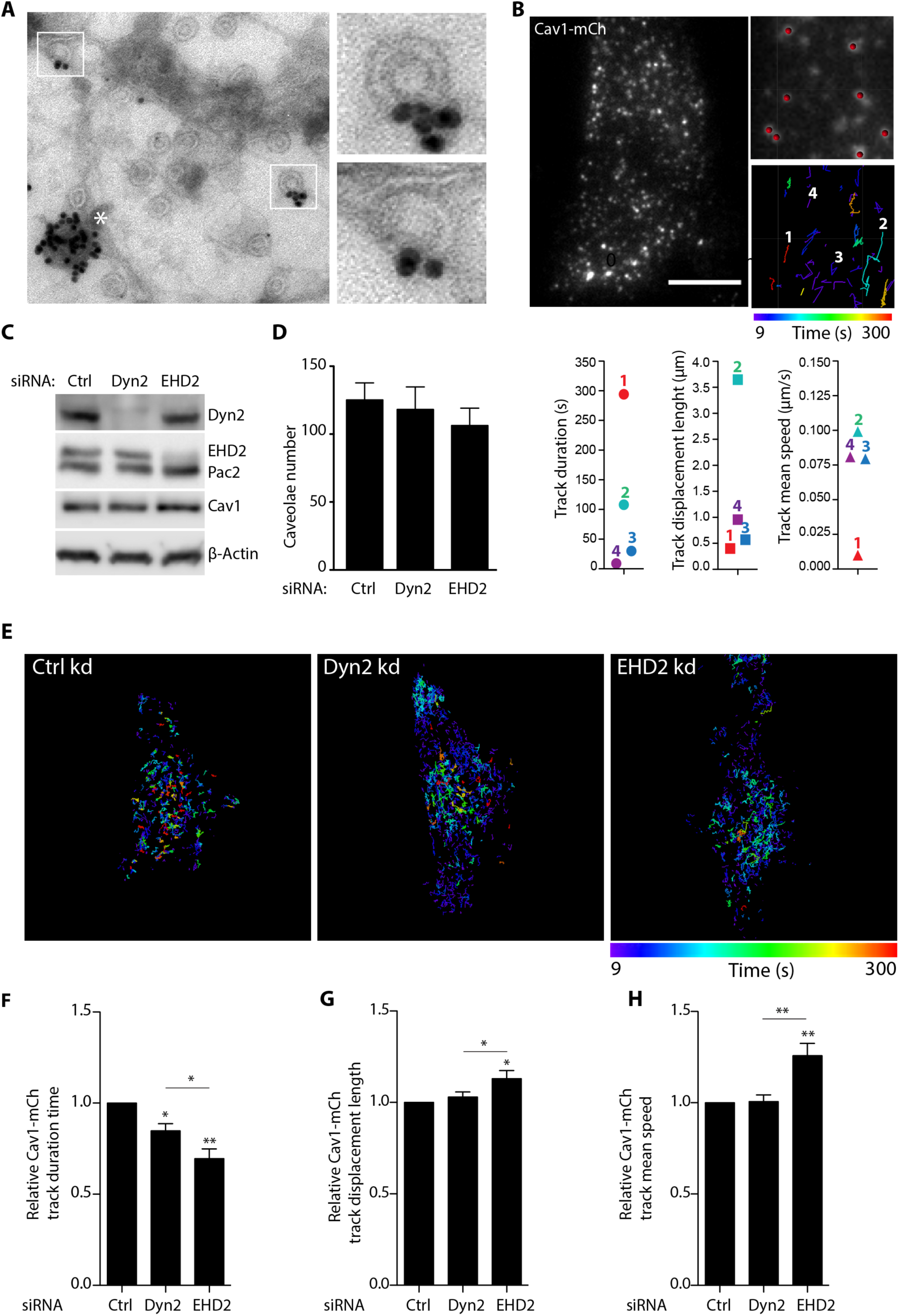
Dynamin2 depletions leads to increased caveolae mobility in Cav1-mCh FlpIn cells. **(A)** Immunogold staining of dynamin2 on plasma membrane lawns prepared from 3T3-L1 cells. (gold particles 15nm). Caveolae in white boxes are shown at higher magnification in right panels; clathrin coated pit marked with asterisk. **(B)** Representative image from TIRF movie of Cav1-mCh cell. Scale bar, 10 μm. Color-coded trajectories illustrate the time and displacement length of tracked caveolae structures during 5 minutes. 1 in red represents caveolae with long duration time, short displacement length and low speed. 2 in cyan represents caveolae with long duration time, short displacement length and low speed. 3 in blue represents caveolae with short duration time, short displacement length and high speed. 4 in purple represents caveolae with short duration time, intermediate displacement length and high speed. (C-H) Cav1-mCh cells were transfected with siRNA for 72 hours before experiments were performed. **(C)** Representative immunoblots of cells treated with siRNAs as indicated. n=3 **(D)** Number of caveolae counted in the basal plasma membrane in cells imaged on TIRF. Mean ± SEM from at least 17 cells per condition. **(E)** Color-coded trajectories of caveolae in Cav1-mCh cells treated with siRNA as indicated. Cells were imaged by TIRF over 5 minutes. **(F)** Quantification of Cav1-mCh track duration time **(G)** speed and **(H)** displacement length. Numbers were related to control (Ctrl) treated cells. Analyses were performed using Imaris software and track mean ± SEM from at least 8 cells per condition are shown. Significance was assessed using *t* test, * p ≤0.05, ** p ≤0.01, *** p ≤0.001.

To determine the role of Dyn2 in regulating caveola dynamics, cells were depleted of Dyn2 using siRNA (Fig. 1C) prior to TIRF microscopy analysis. For comparison, cells were also treated with control siRNA or siRNA against EHD2. Quantification of the number of caveolae at the basal membrane did not reveal any significant change in cells depleted of dynamin or EHD2 as compared to control (ctrl) treated cells (Fig. 1D). Analysis of the dynamic track parameters confirmed that EHD2 depletion, in accordance to its role in stabilizing caveolae at the PM (Moren et al., 2012; Stoeber et al., 2012), resulted in less stable and more fissioned caveolae as compared to ctrl treated cells (Fig. 1E-H and video 2). Surprisingly, dynamin knock down resulted in a 20 percent overall decrease in the duration time, showing that similar to EHD2 depletion, loss of Dyn2 decreased the time that caveolae spent at the PM (Fig. 1E-H and video 3). However, in contrast to EHD2-depleted cells, there was only a slight difference in the speed or displacement as compared to control cells. The uptake of transferrin was severely impaired in Dyn2 depleted cells in agreement with that CCV fission was blocked (Fig. S1). Although low levels of Dyn2 could be remaining in these cells following siRNA depletion, these data suggested that Dyn2 is not needed for caveola fission, but that it rather stabilizes the PM association of caveolae.

### Dyn2 confines caveolae to the PM

To follow up on the potential stabilizing role of Dyn2 at caveolae and enable visualization in real time, we constructed a HeLa FlpIn cell line where the expression of Dyn2-EGFP (Dyn2-GFP) and Cav1-mCherry (Cav1-mCh) could be induced to sub or near endogenous levels (Fig. 2A). TIRF time lapse movies of the cells showed that the majority of the GFP-labeled Dyn2 did not colocalize with caveolae as expected due to its recruitment to clathrin coated vesicles (Fig. 2B). In addition to caveolae only positive for Cav1-mCh, we detected a distinct pool of caveolae where Dyn2-GFP colocalized with Cav1-mCh (Fig. 2B) in accordance with previous result (Fig. 1A). In these cells, which express elevated levels of Dyn2 quantification suggested that this fraction represented approximately 40% of the caveolae (Fig. 2C). The distinct colocalization to a subset of caveolae was confirmed by super resolution SIM microscopy (Fig. S2A and B and video 4). Kymographic analysis of the colocalization over time showed that Dyn2 was stably associated with long-lived caveolae (Fig. 2D and Fig. S2C). Furthermore, the fluorescent intensity profile of Dyn2-GFP and Cav1-mCh showed correlated fluctuations indicative of stable levels of Dyn2-GFP at caveolae over time (Fig. 2E). These results show that the mere presence of Dyn2 at caveolae does not lead to fission.

**Fig. 2.**
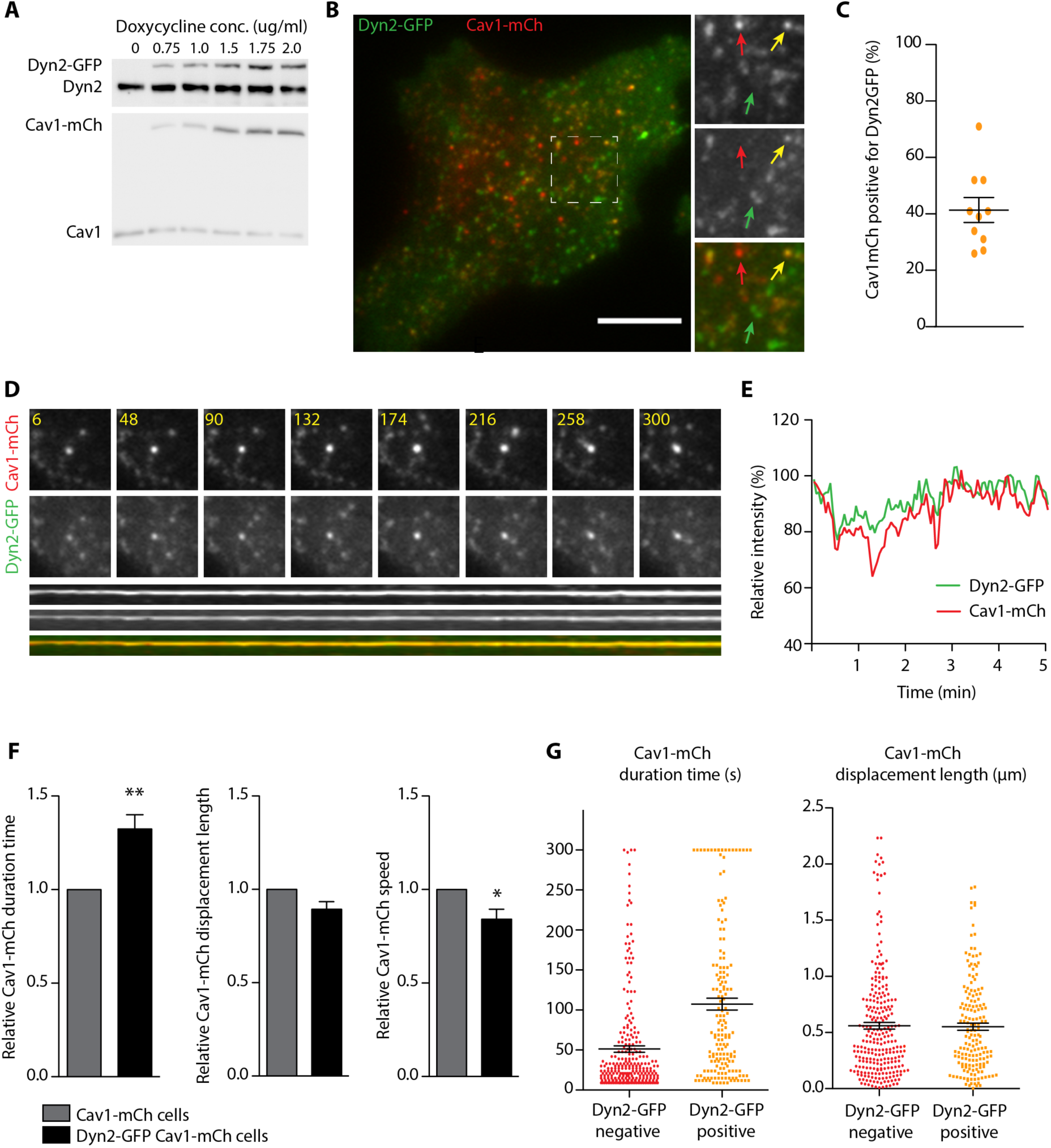
(A) Dyn2-GFP restricts the caveolae mobility in the PM. Representative immunoblot of Dyn2-GFP-Cav1-mCh cells treated with concentrations of doxycycline as indicated. Doxycycline concentration used for further experiments were 1.0 ng/ml. **(B)** Representative image from TIRF movie of Dyn2-GFP-Cav1-mCh cell. Red arrow highlight structure only positive for Cav1-mCh, yellow arrow highlight structure positive for both Cav1-mCh and Dyn2-GFP and green arrow highlight structures only positive for Dyn2-GFP. Scale bar, 10 μm. **(C)** Quantification of percentage of caveolae positive for Dyn2-GFP. Data are shown as scatter dot plot, mean ± SEM. **(D)** Top, TIRF image series of stable caveolae positive for both Cav1-mCh and Dyn2-GFP. Time in sec as indicated. Bottom, kymograph showing stable association of Cav1-mCh and Dyn2-GFP to the plasma membrane during 300 sec in TIRF movie. **(E)** Relative fluorescence intensity of mCh and GFP fluorophore of caveolae positive for both Cav1-mCh and Dyn2-GFP over time. **(F)** Quantification of Cav1-mCh track duration time, displacement length and mean speed in Cav1-mCh cells (grey) and Dyn2-GFP-Cav1-mCh cells (black). Analysis were performed using Imaris software and track mean from at least 10 cells per condition are shown ± SEM. **(G)** Caveolae track duration times and displacement lengths divided in pools of Dyn2-GFP negative (red) or positive (orange). Dyn2-GFP-Cav1-mCh cells were imaged on TIRF over 5 minutes. Cav1-mCh spots were followed throughout the time series and scored whether they were positive or negative for Dyn2-GFP. Scatter dot plots show all Cav1-mCh tracks from three different cells. Analysis were performed using Imaris software and data are shown as mean ± SEM. Significance was assessed using *t* test, * p ≤0.05, ** p ≤0.01.

To study if the elevated levels of dynamin affected the time caveolae spent at the cell surface Cav1-mCh spots were tracked and the dynamic parameters were analyzed (Fig. 2F). We found that the track duration time in cells expressing Dyn2-GFP increased by around 30% as compared to caveolae tracks in Cav1-mCh FlpIn cells (Fig. 2F). Furthermore, we observed a decrease in the displacement length and mean speed (Fig. 2F), further supporting that Dyn2 stabilizes caveolae and prevent fission. As elevated global levels of Dyn2 might have an overall effect on the cell, and thereby indirectly increase the caveolae stability, we aimed to determine if the stabilizing effect was specific and directly coupled to the localization of Dyn2 at caveolae. Caveolae were divided into two pools, positive or negative for Dyn2-GFP, and the track duration time for each pool was measured and plotted (Fig. 2G). Caveolae positive for Dyn2-GFP displayed a mean duration time that was more than twice as long as the pool that lacked Dyn2-GFP, showing that Dyn2 stabilized caveolae at the PM. Interestingly, the displacement length of the different pools was not significantly different in these cells as compared to control (Fig. 2G). This agrees with the finding that knock down (KD) of Dyn2 did not have a significant effect on the displacement length and suggests that Dyn2 regulates cell surface duration of caveolae without influencing the lateral movement.

### Dyn2 recruitment to caveolae is independent of EHD2 and pacsin2

As Dyn2, similarly to EHD2, confined caveolae to the cell surface, we wanted to compare the localization and function of these proteins at caveolae. Immunogold co-labeling of Dyn2 and EHD2 in 3T3-L1 cells showed that EHD2 was present at most caveolae, while Dyn2 labeling was only observed on a subset. Importantly, caveolae positive for Dyn2 were also positive for EHD2 showing that recruitment of these proteins is not mutually exclusive (Fig. 3A). To monitor caveolae recruitment in live cells, BFP-tagged EHD2 was overexpressed in induced Dyn-GFP-Cav1-mCh HeLa FlpIn cells followed by TIRF imaging (Fig. 3B). We found that Dyn2 indeed localized together with EHD2 on the same caveolae. Dyn2 spots devoid of Cav1-mCh were always free from EHD2-BFP (Fig. 3B). Dynamin has been suggested to interact with EHDs and pacsins (Jakobsson et al., 2011; Qualmann et al., 1999). In order to test if Dyn2 was dependent on EHD2 or pacsin2 for its caveola-recruitment, these proteins were depleted in Dyn2-GFP-Cav1-mCh FlpIn cells and imaged using TIRF. Dyn2-GFP localized to Cav1-mCh spots regardless of whether EHD2 or pacsin2 were depleted or not (Fig. S3A). In addition, the fluorescent recovery after photobleaching revealed no change in recovery rate of Dyn2-GFP at caveolae in the absence of EHD2 or pacsin2 as compared to ctrl cells (Fig. 3C). These results show that Dyn2 is not dependent on EHD2 nor pacsin2 for its recruitment to caveolae.

**Fig. 3.**
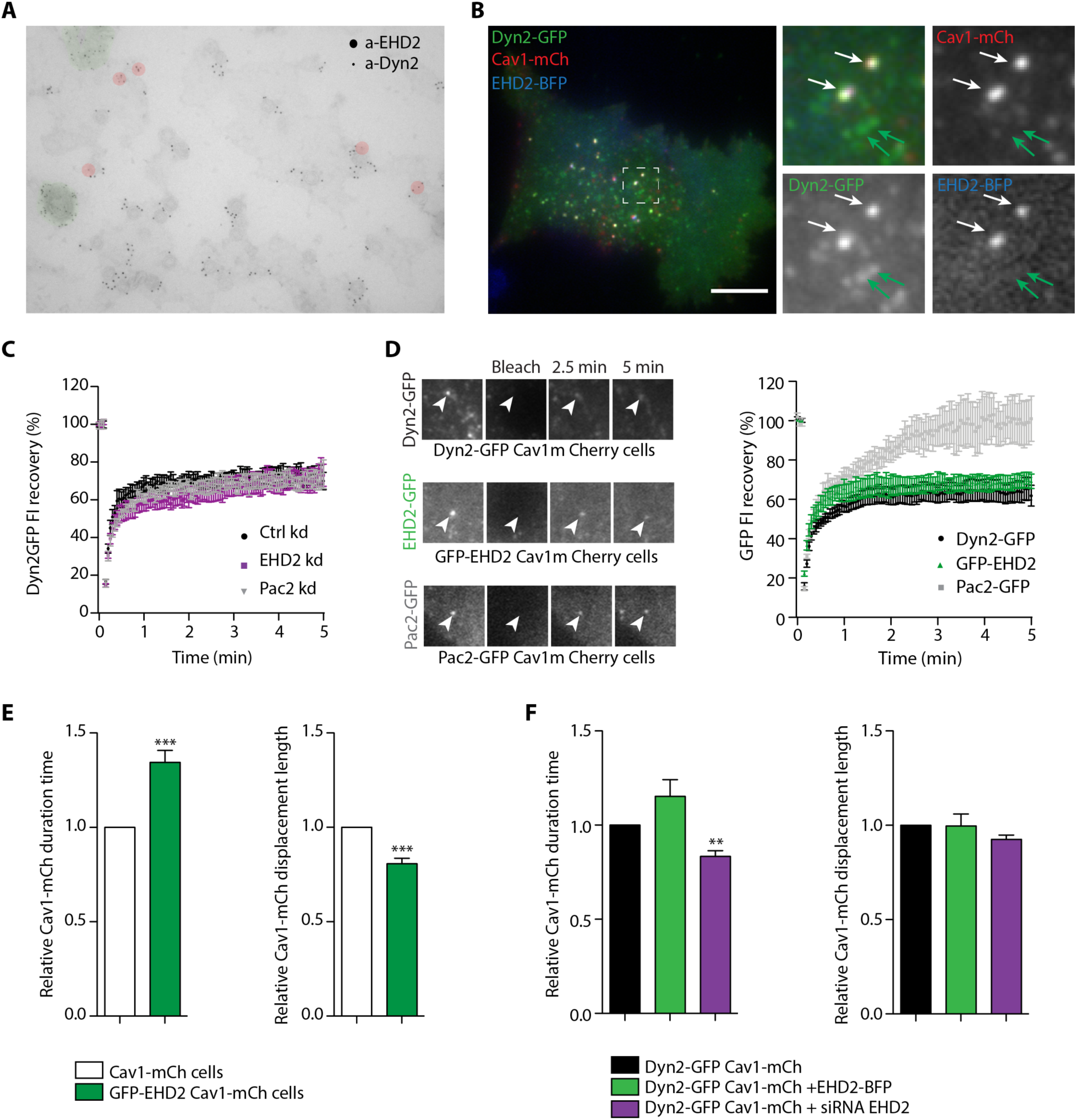
Localization of Dyn2-GFP to caveolae is independent of EHD2 and pacsin2 presence. **(A)** Immunogold labelling of EHD2 (large gold) and Dyn2 (small gold) on plasma membrane lawns prepared from 3T3-L1 cells. Red indicates caveolae and green clathrin coated pits. **(B)** Representative image from TIRF movie of Dyn2-GFP-Cav1-mCh cells transiently expressing EHD2-BFP. White arrows highlight structures where all three tagged proteins are colocalized and green arrows highlights structures only positive for Dyn2-GFP. Scale bar, 10 μm. **(C)** Recovery curves of Dyn2-GFP intensities after photobleaching of caveolae in Dyn2-GFP-Cav1-mCh cells treated with Ctrl, EHD2 or pacsin2 siRNA as indicated. **(D)** Image panel showing GFP recovery after photobleaching in Dyn2GFP-Cav1-mCh cells, GFP-EHD2-Cav1-mCh cells or Pac2-GFP-Cav1-mCh cells. Graph shows percentage GFP-intensity recovery curves after photobleaching of mCh and GFP colocalizing structures at the PM. **(E)** Quantification of Cav1-mCh track duration time and displacement length in GFP-EHD2-Cav1-mCh cells (green). Numbers were related to Cav1-mCh cells (white). Track mean ± SEM from at least 7 cells per condition are shown. **(F)** Quantification of Cav1-mCh track duration time and displacement length in Dyn2-GFP-Cav1-mCh cells (black), transiently expressing EHD2-BFP (dark green) or depleted from EHD2 (purple). Numbers were related to Dyn2-GFP-Cav1-mCh cells (black). Track mean ± SEM from at least 7 cells per condition are shown. Significance was assessed using *t* test, ** p ≤0.01, *** p ≤0.001.

### Dyn2 and EHD2 cooperatively stabilize caveolae to the PM

To be able to compare the role of Dyn2, EHD2 and Pacsin2 (Pac2) at caveolae, we constructed GFP-EHD2-Cav1-mCh and pacsin2-Cav1-mCh HeLa FlpIn cell lines (Fig. S3B and C). GFP-EHD2 was detected on most caveolae in agreement with previous results (Fig. S3B) (Moren et al., 2012; Stoeber et al., 2012). Pac2-GFP was observed to tubulate membrane devoid of Cav1-mCh, but also to colocalize to Cav1-mCh, however not to the same extent as Dyn2 or EHD2 (Fig. S3C). To compare the turnover rates at caveolae, Dyn2-GFP, GFP-EHD2 or GFP-Pac2 were photobleached at caveolae in the respective cell lines, and the fluorescence recovery was recorded and plotted (Fig. 3D). All three proteins showed a similar initial rate of recovery following photobleaching. However, five minutes after photobleaching only approximately 60 percent of the fluorescent signal had recovered for both Dyn2 and EHD2 (Fig. 3D). The low recovery showed that there is a large, stably associated pool of both Dyn2 and EHD2 at caveolae. In comparison, the entire pool of fluorescent pacsin2 had been exchanged after five minutes (Fig. 3D). This shows that dynamin2 and EHD2 are firmly associated with caveolae.

Our data shows that both Dyn2 and EHD2 stabilize caveolae. Yet, Dyn2 is only localized to a subset of stabile caveolae, implying that Dyn2 and EHD2 might stabilize caveolae via different but cooperative mechanisms. When comparing the surface duration of caveolae in EHD2-Cav1-mCh HeLa FlpIn cell lines we noted that similar to Dyn2, the extra EHD2 aids in caveolae stabilization (Fig. 3E). Interestingly, however, while caveola displacement was not altered in Dyn2-Cav1-mCh cells, the displacement was significantly shorter in EHD2-Cav1-mCh cells as compared to control cells (Fig. 3E and Fig. S3D). This shows that EHD2 but not Dyn2 influence the lateral movement of caveolae. To further determine if Dyn2 and EHD2 could act cooperatively to stabilize caveolae, we tracked and compared the dynamic track parameters in Dyn-GFP-Cav1-mCh HeLa FlpIn cells following expression of BFP-tagged EHD2. Analysis showed that overexpression of BFP-EHD2 further increased the duration time of caveolae in relation to the already stabilizing effect of Dyn2 in these cells. In agreement with that Dyn2 and EHD2 cooperatively affects caveolae stability, depletion of EHD2 from the Dyn2-GFP-Cav1-mCh cells decreased the caveolae duration time, but not to the level of EHD2-depleted Cav1-mCh cells (Fig. 3F and 1F). This showed that Dyn2 still has a stabilizing effect even following EHD2 depletion suggesting that the proteins might act via distinct mechanisms.

### Dyn2 interacts with Cav1 at the caveolae bulb

Our data suggests that EHD2 and Dyn2 work independently at caveolae, yet both proteins are known to oligomerize into rings and spirals on highly curved membranes such as the caveolae neck. To decipher whether Dyn2 and EHD2 both were located at the caveolae neck, we expressed the mutant ΔNΔEH-EHD2, known to extend the caveolae neck into long tubules (Hoernke et al., 2017), in the in Dyn2-GFP-Cav1-mCh HeLa FlpIn cells (Fig. 4A). Tubes decorated by BFP-EHD2 were lacking Dyn2-GFP, which instead colocalized with Cav1-mCh in puncta at the end of the tubes (Fig. 4A), suggesting that Dyn2 localizes to the bulb. Consistent with this, immunogold labelling also showed no preferential localization to the caveolar neck under our labelling conditions (Fig. 1A).

**Fig. 4.**
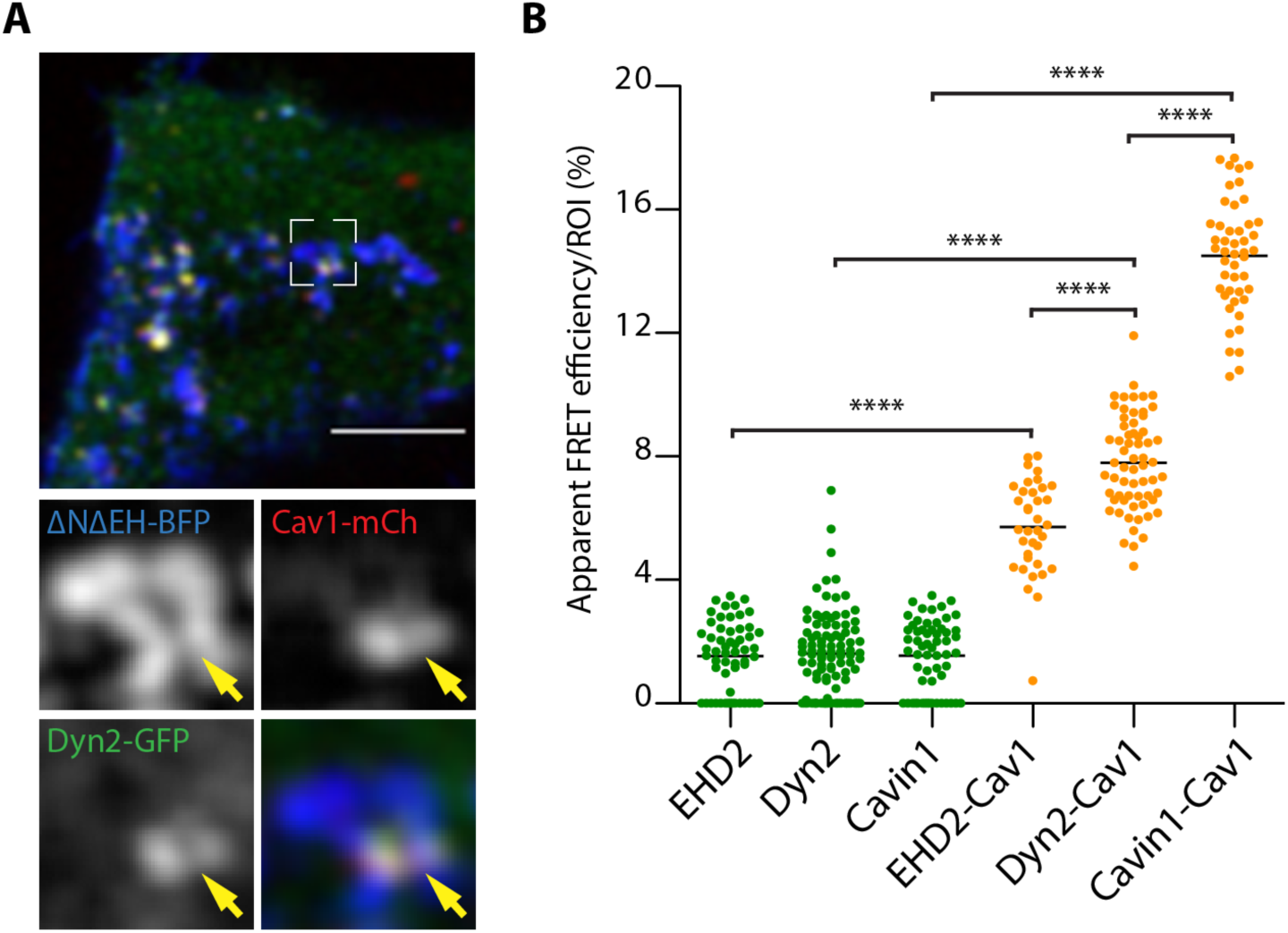
Dyn2-GFP localizes closer to the caveolae bulb than EHD2. **(A)** Confocal image of a Dyn2-GFP-Cav1-mCh cell transiently expressing Δ NΔEH EHD2-BFP. Yellow arrow highlights Dyn2-GFP localization to Cav1-mCh at the tip of the membrane tube. Scale bar, 10 μm. **(B)** Apparent FRET efficiency between GFP and mCh fluorophores in GFP-EHD2-Cav1-mCh cells, Dyn2-GFP-Cav1-mCh cells or Cav1-mCh transiently expressing Cavin1-GFP. ROIs of structures positive for only GFP (green) or GFP and Cav1 (orange) were used to calculate the apparent FRET efficiency. Significance was assessed using *t* test, **** p ≤0.0001.

To further address the interactions of caveolae components in live cells, we used fluorescence life time microscopy (FLIM). This methodology is based on FRET energy transfer between fluorophores in close proximity affecting the fluorescence life time. First, the life time of singly expressed Dyn2-GFP, GFP-EHD2 or Cavin1-GFP was determined (Fig. 4B and Fig. S4). Next, FLIM was measured on structures where Cav1-mCh colocalized together with either Dyn2-GFP, GFP-EHD2 or Cavin1-GFP and the apparent FRET efficiency was calculated (Fig. 4B). Cavin1, which together with caveolin1 builds up the caveolae coat, resulted in a very high FRET efficiency. This shows that, as expected, Cavin1 and Cav1 interact in cells. However, EHD2, which localizes at the caveolae neck, showed a low degree of FRET efficiency (Fig. 4B). Interestingly, the FRET efficiency for dynamin was intermediate (Fig. 4B). These data suggest that dynamin and EHD2 are not situated at the same position on caveolae but rather that dynamin localizes closer to the caveolae bulb.

### Dyn2 GTPase activity is not required for sequestering of caveolae to the PM

Given that dynamin, independently of EHD2, stabilized caveolae we aimed to investigate if the GTP cycle of Dyn2 influenced its role at caveolae. As a means to manipulate the intrinsic activity of dynamin we employed the commonly used transition-state-deficient dynamin mutant K44A. This mutant has been shown to oligomerize, but to have decreased GTP affinity and hydrolysis rate, and its expression in cells inhibit fission of clathrin coated vesicles (Damke et al., 1994). It should be noted that transiently overexpressed Dyn2 as well as Dyn2 K44A mislocalizes to stable aggregates (Fig. S5A). Therefore, we constructed inducible K44A-Dyn2-GFP-Cav1-mCh HeLa FlpIn cells (Fig. 5A and S5A). Cells imaged by TIRF revealed that Dyn2 K44A also localized to caveolae, although the percentage of Cav1-mCh spots colocalizing with dynamin was lower than that of wild type dynamin (Fig. 5A and B). This is likely due to the impaired GTP binding and hence lower membrane recruitment of K44A. Therefore, to be able to compare the effects on caveolae stability, caveolae were divided into pools positive versus negative for K44A Dyn2-GFP. Tracking of the caveolae dynamics showed that the presence of Dyn2 K44A increased the Cav1-mCh track duration time to a similar extent as to that of wild type dynamin (Fig. 5C and Fig. 2G). This showed that GTP hydrolysis was not required for the ability of Dyn2 to stabilize caveolae at the plasma membrane. Similar to wild type Dyn2, expression of K44A can still be used to restrict caveolae fission, as previously suggested, but not due to its GTPase deficiency but rather since it able to form oligomers at caveolae which is sufficient for caveolae stabilization.

**Fig. 5.**
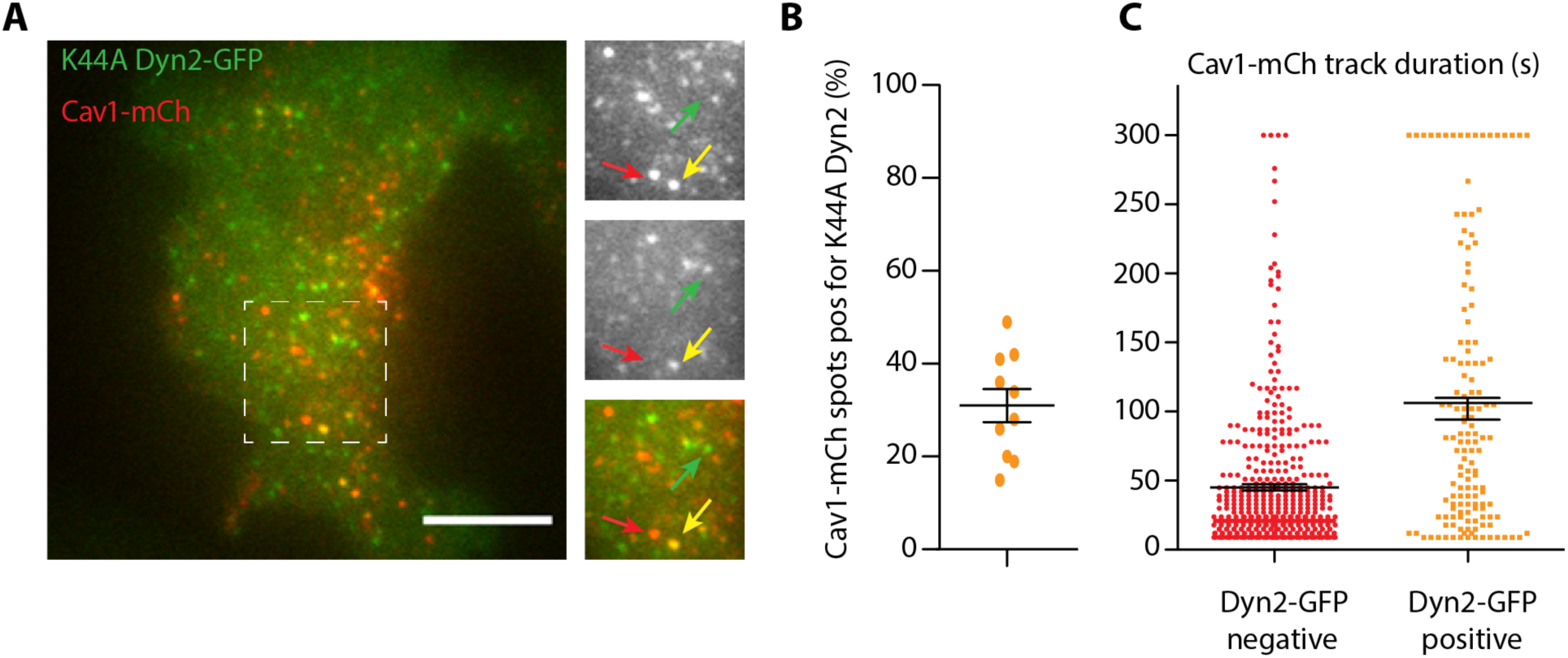
K44A Dyn2-GFP localizes to and stabilize caveolae to the PM. **(A)** Representative image from TIRF movie of K44A Dyn2-GFP-Cav1-mCh cell. Red arrow highlight structure only positive for Cav1-mCh, yellow arrow highlight structure positive for both Cav1-mCh and K44A Dyn2-GFP and green arrow highlight structures only positive for K44A Dyn2-GFP. Scale bar, 10 μm. **(B)** Quantification of percentage of caveolae positive for K44A Dyn2-GFP. Data are shown as scatter dot plot, mean ± SEM. **(C)** Caveolae track duration times and displacement lengths divided in pools of K44A Dyn2-GFP negative (red) or positive (orange). K44A Dyn2-GFP-Cav1-mCh cells were imaged on TIRF over 5 minutes. Cav1-mCh spots were followed and scored whether they were positive or negative for Dyn2-GFP. Graph shows all Cav1-mCh tracks from three different cells. Analysis were performed using Imaris software and data are presented as scatter dot plot, mean ± SEM. Significance was assessed using *t* test.

### Dyngo4a immobilize caveolae at the plasma membrane

The small molecule dynamin inhibitor Dyngo 4a has been shown to prevent membrane-stimulated GTPase activity of dynamin and halt CCV endocytosis (McCluskey et al., 2013). Since the GTPase deficient mutant K44A was able to stabilize caveolae, we wanted investigate the effect of Dyngo 4a on caveola dynamics. Dyngo 4a was added to Dyn-GFP-Cav1-mCh HeLa FlpIn cells and incubated for 30 minutes followed by TIRF imaging. As previously demonstrated cells retracted in response to Dyngo 4a treatment (Park et al., 2013), reducing the basal surface area (Fig S6A). Indeed, Dyngo 4a was reported to have off target effects on fluid-phase endocytosis and the actin cytoskeleton and we could confirm that the F-actin was disrupted in cells treated with Dyngo 4a as compared to mock-treated cells (Fig. S6B). The number of Cav1-mCh spots at the basal surface was also dramatically reduced (Fig. 6A and B), but, 70 percent of the remaining spots were positive for Dyn2 (Fig. S6C). We could confirm that the spots were surface connected caveolae as they were positive for Cavin1 and EHD2 (Fig. S6D). The increased ratio of caveolae positive for Dyn2 could suggest that these caveolae were more resistant to Dyngo 4a treatment, or that the inhibition of the GTPase activity stabilized more Dyn2 oligomers at caveolae. As previously shown, CCV-mediated uptake of transferrin was blocked in Dyngo 4a treated cells (Fig. S6E).

**Fig. 6.**
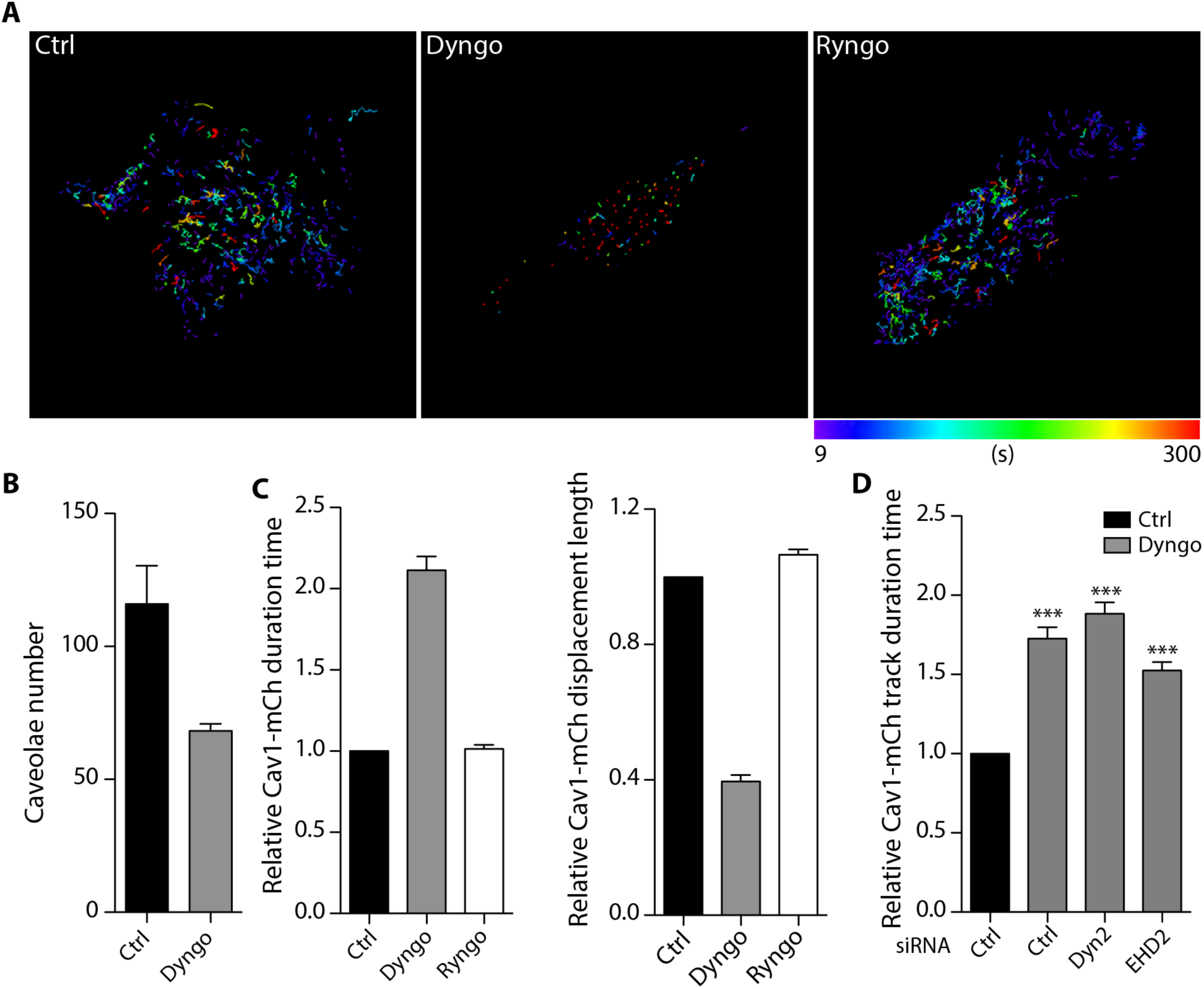
Dyngo 4a stabilization of caveolae is independent of dynamin involvement. **(A)** Color-coded trajectories of caveolae in Dyn2-GFP-Cav1-mCh cells pretreated with DMSO (Ctrl), Dyngo 4a or Ryngo 1-23 for 30 minutes before imaged on TIRF over 5 minutes. **(B)** Quantification of percentage of caveolae positive for Dyn2-GFP. Data are shown as scatter dot plot, mean ± SEM. **(C)** Quantification of Cav1-mCh track duration time and displacement length in cells as indicated. Numbers were related to control (Ctrl) treated cells. Analysis were performed using Imaris software and track mean from at least 8 cells per condition are shown ± SEM. **(D)** Quantification of Cav1-mCh track duration time in Cav1-mCh cells treated with siRNA as indicated prior to pretreatment of DMSO (Ctrl) or Dyngo 4a as in (A). Numbers were related to DMSO-treated Ctrl siRNA-treated cells (black bar). Analysis were performed using Imaris software and track mean from at least 8 cells per condition are shown ± SEM. Significance was assessed using *t* test, *** p ≤0.001.

Tracking of Cav1-mCh spots in Dyngo 4a treated cells revealed a dramatic effect on the caveola dynamics (Fig. 6A and C and video 5). The track duration times were increased to the double and the track displacement length had decreased to less than half of that of ctrl treated cells (Fig. 6A and C). This tracking pattern revealed that caveolae had become almost static in comparison to mock-treated cells (Fig. 6A). This was surprising since these effects on caveola dynamics were several times higher than the stabilizing effect of Dyn2 or Dyn2 K44A or the increased mobility observed in cells depleted of Dyn2 or EHD2. To determine if the dramatic effects were indeed due to inhibition of dynamin GTPase activity, the Dyngo 4a treatment was repeated in Cav1-mCh FlpIn HeLa cells depleted of Dyn2. Surprisingly, the drug had the same stabilizing effect on the caveolae in these cells (Fig. 6D). Similarly, EHD2 KD did not significantly influence the stabilizing effect of Dyngo 4a. These data showed that caveolae can indeed be immobilized, using Dyngo 4a as previously proposed, but that this effect is not due to block of Dyn2 GTPase activity.

In view of these results, which go against a long-standing dogma that has developed in the field, we wanted to test if stimulation of the actin-dependent oligomerization of Dyn2 into rings influenced its stabilizing effect on caveola dynamics. For this we used Ryngo 1-23, a stimulator of dynamin GTPase activity by actin-dependent oligomerization (Gu et al., 2014). This dynamin modulator has been shown to induce Dyn2-dependent actin filament rearrangements (Lees et al., 2015), and increase dynamin-dependent stabilization of the fusion pore during exocytosis (Jackson et al., 2015). As previously shown (Sandgren et al., 2010), Ryngo 1-23 treatment impaired the uptake of transferrin in cells (Fig S6E). Next, we analyzed its effect on caveolae by TIRF analysis and tracking of Cav1-mCh (Fig. 6A and C). Analysis of the dynamic track parameters showed that the

Ryngo 1-23-induced increase of dynamin GTPase activity had no apparent effect on the caveolae track duration time, or displacement length at the PM as compared to ctrl treated cells (Fig. 6A and C). Taken together, our results show that stimulation or inhibition of the GTPase activity of Dyn2 does not influence caveola dynamics, but that Dyngo 4a has dramatic, Dyn2 independent, effects on caveola stability.

### Lipid induced fission of caveolae is dynamin independent

Increased levels of cholesterol in the plasma membrane has been shown to drive fission of caveolae (Hubert et al., 2020). To address if Dyn2 was involved in this process, we used the same methodology to rapidly incorporate cholesterol in the plasma membrane (Fig. 7A and B). Real time TIRF imaging showed that the plasma membrane levels of fluorescent cholesterol indeed increased over time (Fig. 7B). Cav1-mCh cells transfected with ctrl or Dyn2 siRNA were treated with cholesterol-containing fusogenic liposomes for 15 minutes before being imaged on TIRF. As expected, tracking analysis showed that cholesterol-addition to Cav1-mCh cells significantly decreased the time that the caveolae spent at the PM compared to non-treated cells (Fig. 7C). Interestingly, the duration time in cells depleted of Dyn2 was also reduced by cholesterol addition and to the same level as in cells treated with ctrl siRNA (Fig. 7C). This revealed that Dyn2 was not required for cholesterol driven fission of caveolae.

**Fig. 7.**
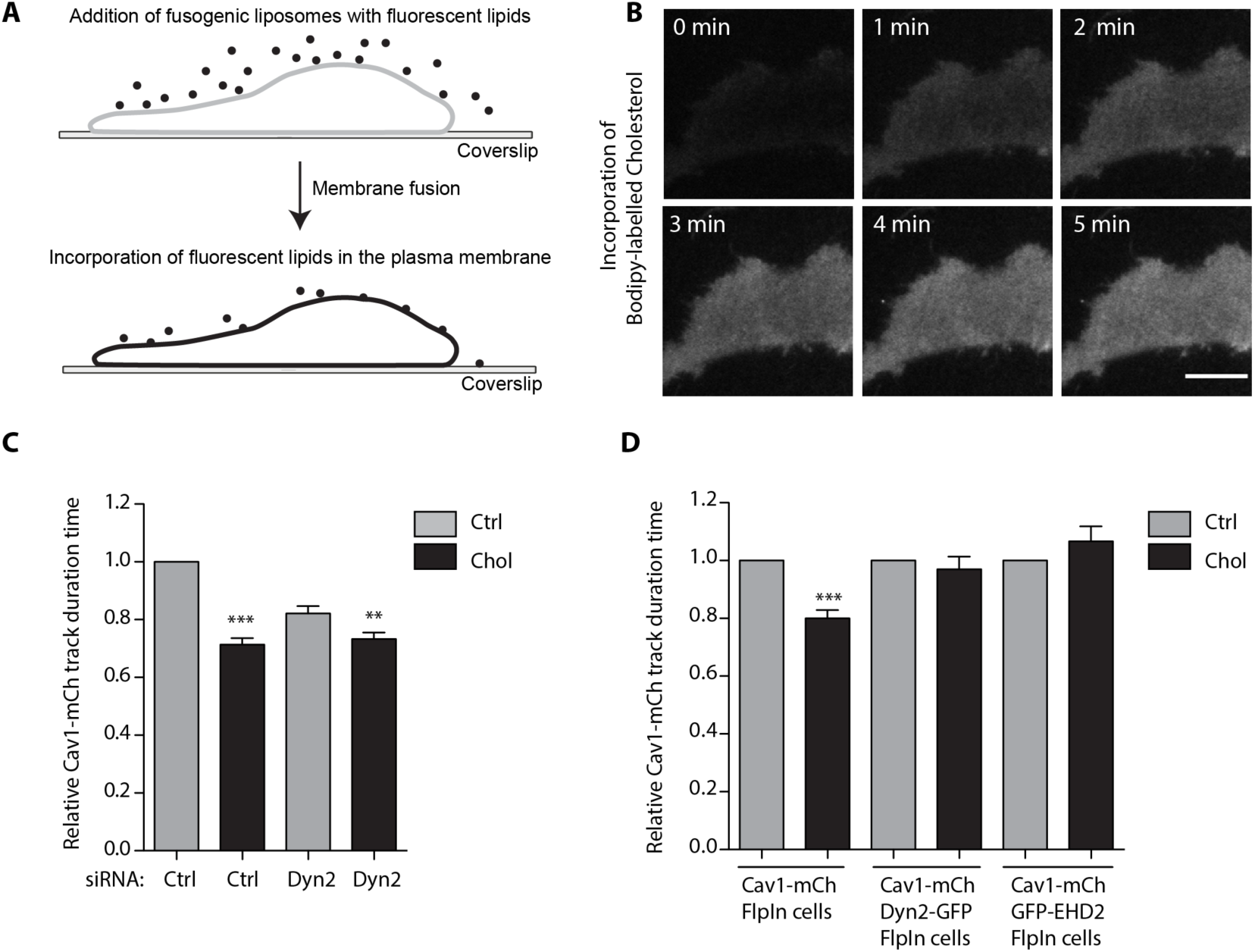
Lipid induced fission of caveolae is counteracted by Dyn2-GFP. **(A)** Schematic of fusogenic liposome methodology. **(B)** Time series showing the incorporation of Bodipy-labelled cholesterol into the PM of a Cav1-mCh cell. Scale bar, 10 μm. **(C)** Quantification of track duration time in Cav1-mCh cells pretreated with Ctrl or Dyn2 siRNA as indicated. Cells were non-treated (grey bar) or treated with fusogenic liposomes containing cholesterol (black bar) for 20 minutes prior to imaging. Analyses were performed using Imaris software and track mean from at least 10 cells per condition are shown ± SEM. **(D)** Quantification of track duration time in Cav1-mCh cells, Dyn2-GFP-Cav1-mCh cells or GFP-EHD2-Cav1-mCh cells non-treated (grey bar) or treated with fusogenic liposomes containing cholesterol (black bar) for 20 minutes prior to imaging. Analysis were performed using Imaris software and track mean from at least 10 cells per condition are shown ± SEM. Significance was assessed using *t* test, ** p ≤0.01, *** p ≤0.001.

To test if Dyn2 instead could restrain the lipid-driven fission of caveolae, the effects of cholesterol incorporation on caveola dynamics in Dyn2-GFP-Cav1-mCh cells was monitored and compared to the dynamics in Cav1-mCh cells. Interestingly, cholesterol administration had no significant effect on the caveola track duration in Dyn2-GFP-Cav1-mCh cells (Fig. 7D), showing that a slight excess of Dyn2 was enough to restrain the lipid-driven fission. Similarly, the elevated levels of EHD2 in GFP-EHD2-Cav1-mCh FlpIn cells also prevented cholesterol induced fission (Fig. 7D), as previously shown (Hubert et al., 2020). These data show that dynamin, just like EHD2, prevents caveolae-fission induced by cholesterol sequestering and furthermore that lipid-induced fission of caveolae from the PM is dynamin independent. Taken together, our results show that Dyn2 significantly contributes to the atypical dynamics of caveolae by stabilizing the PM association of these specialized membrane domains, supporting a dual role of dynamin oligomerization at membrane neck regions.

## Discussion

All membrane vesicle fission and fusion processes are known to proceed through an intermediate state characterized as a membrane pore or neck transiently connecting the vesicle to the membrane. Yet, caveolae display atypical dynamics in the sense that such invaginations are stably associated with the PM over time. Dynamin-like proteins make up a family of homotypic membrane fission and fusion proteins known to stabilize or destabilize membrane necks and pores via nucleotide dependent oligomerization. Herein we have studied the role of Dyn2 on caveola dynamics in living cells using TIRF microscopy and single particle tracking. In contrast to its role in CCV fission, we find that Dyn2 acts as a stabilizing protein, increasing the time caveolae spends at the PM. Dyn2 was not found on all caveolae, but the Dyn2-positive caveolae displayed PM duration times that were twice as long as the caveolae devoid of Dyn2. Depletion of Dyn2 from cells destabilized the PM association of caveolae leading to shorter duration times. Furthermore, Dyn2 was not required for lipid-induced caveolae fission as triggered by exogenous addition of cholesterol. However, excess levels of Dyn2 could counteract such lipid-induced fission. This stabilizing role of Dyn2 did not require GTP-hydrolysis, but did seem to depend on Dyn2 oligomerization. Based on our data, we find that dynamin is not required for caveola formation or fission. We propose that it acts cooperatively to EHD2, but via an independent mechanisms, to restrain a subset of caveolae to the PM.

Our results, which go against the current dogma on caveolae fission (Matthaeus and Taraska, 2020), can be reconciled with previous data, and build a refined model for how Dyn2 influence caveola dynamics. Dyn2 has been previously shown to localize to caveolae (Oh et al., 1998) in agreement with our findings. Yet, the amount of caveolae which are positive for Dyn2 is difficult to measure and could vary in between cell types and the techniques used. Immunogold labelling of endogenous Dyn2 in 3T3L1 cells suggested that rather a small fraction of caveolae were positive for Dyn2, but this could be underestimating the amount based on poor antibody binding. In the Dyn2-GFP-Cav1-mCh cells, we found that approximately 40 percent of the caveolae were positive for dynamin, but these cells have elevated levels of Dyn2. However, taken together, all data support that a distinct pool of caveolae are positive for Dyn2. Dynamin and GTP-hydrolysis was first proposed to potentiate fission of caveolae based on release of Cav1 from purified plasma membranes following incubation with cytosol and GTP (Schnitzer et al., 1995; Schnitzer et al., 1996). Additionally, microinjection of antibodies directed against Dyn2 in hepatocytes greatly decreased the uptake of Cholera toxin, which was interpreted to indicate that the internalization of caveolae had been blocked (Henley et al., 1998). However, Cav1 has been shown to be released from the PM by alternative vesicles in the absence of the caveolae coat machinery, and cholera toxin has been shown to internalize via other clathrin-independent mechanisms as reviewed in (Kenworthy et al., 2021). This demonstrates the need for direct analysis of caveola dynamics in living cells. Here, we have used inducible cell lines with expression levels of caveolin and dynamin close to endogenous levels in combination with TIRF microscopy and single particle tracking of caveolae to determine their PM duration, lateral displacement and speed on a large number of caveolae under different conditions.

In addition to Dyn2, caveolae stabilization has been shown to depend on peripheral membrane proteins such as EHD2 and pacsin2. Although dynamins has been suggested to interact with both EHDs and pacsins (Jakobsson et al., 2011; Qualmann et al., 1999), we found that Dyn2 is recruited to caveolae independently of EHD2 and pacsin2. We show that while EHD2 seem to be present at the majority of caveolae, Dyn2 and pacsin2 is limited to a subset. Yet, immunogold labeling and TIRF microscopy revealed that Dyn2 and EHD2 could be simultaneously detected on the same caveolae. By extending the caveolae neck into tubes using the ΔNΔEH-EHD2 mutant we found that Dyn2 was not enriched at such tubules but rather localized closer to the bulb together with Cav1. FLIM-FRET analysis verified that Dyn2 was in closer proximity to Cav1 than what EHD2 was. These data are in agreement with previous results suggesting that Dyn2 and Cav1 interact directly (Yao et al., 2005). Furthermore, the detection of a sub-stochiometric interaction between Dyn2 and Cav1-vesicles (10-15 Dyn2 molecules per 100-150 Cav1) in a cell free system further supports the direct interaction at the caveolae bulb (Jung et al., 2018).

Although both Dyn2 and EHD2 stabilize caveolae, our data suggest that they act via different mechanisms. Analysis showed that overexpression of BFP-EHD2 in Dyn2-GFP-Cav1-mCh cells further increased the duration time of caveolae in relation to the already stabilizing effect of Dyn2 in these cells. Similarly, depletion of EHD2 from the Dyn2-GFP-Cav1-mCh cells decreased the caveolae duration time, but not to the level of Cav1-mCh cells (Fig. 3F and 1F). Furthermore, depletion of Dyn2 or EHD2 in Cav1-mCh cells both resulted in decreased caveola duration time. However, while depletion of EHD2 led to longer caveolae displacement length, this parameter was unaffected in Dyn2 depleted cells. Likewise, increasing the levels of Dyn2 or EHD2 in the inducible cell lines resulted in caveola stabilization. However, the shorter displacement length found in GFP-EHD2-Cav1-mCh expressing cells could not be observed in Dyn2-GFP-Cav1-mCh cells. Displacement of caveolae in the plane of the membrane has been shown to depend on the F-actin (Thomsen et al., 2002), and caveolae are frequently found to align with actin filaments. Coupling of caveolae to F-actin via Filamin A (Stahlhut and van Deurs, 2000), an actin binding protein, was found to increase internal trafficking of caveolae (Sverdlov et al., 2009). Moreover, both EHD2 and Dyn2 have been suggested to interact with actin and to regulate the actin polymerization (Gu et al., 2010; Zhang et al., 2020). It is possible that both Dyn2 and EHD2 form nucleotide dependent oligomers at caveolae which stabilize the membrane structure, but that they influence actin coupling and polymerization differently which affects caveolae displacement in the membrane.

Our data clearly show that by just slightly increasing the cellular levels of Dyn2, caveolae were stabilized and that Dyn2 firmly associated with caveolae over time. This pattern of Dyn2 localization differed to that seen for CCVs, where a spike in Dyn2 recruitment is followed by CCV fission (Merrifield et al., 2002). Furthermore, FRAP analysis showed that a large pool of Dyn2, similar to EHD2, is sequestered at caveolae suggesting that Dyn2 oligomers are formed at caveolae and that GTPase-triggered disassembly of Dyn2 is slow. This also implied that GTP-hydrolysis might not be required for stabilization of caveolae at the PM. In agreement with this, the GTP hydrolysis mutant K44A stabilized PM association of caveolae similar to that seen with wild type Dyn2, although this mutant inhibits fission of CCV. This showed that GTP-hydrolysis by Dyn2 in fact is not required for caveola stabilization. This is consistent with the observation that GTP-hydrolysis facilitates membrane tubule constriction. Dyn2K44A is however known to still form oligomers which are trapped in a transition state-like conformation, which appear to be sufficient for caveolae stabilization. Dynamins have been clearly shown to affect the fusion pore closure. Interestingly, Dyn1 has been described to bi-directionally regulate transmitter release from exocytic vesicles by acting on fusion pore expansion. This suggests that dynamin oligomers might be able to stabilize and perform fission of membrane necks depending on the GTPase cycle (Lasic et al., 2017). Indeed, by the use of the dynamin inhibitor Dyngo 4a, the authors showed that inhibition of GTPase activity reduced vesicular release. However, stimulation of GTPase activity via the modulator Ryngo 1-23 prolonged the time for full fusion and thereby reduced the number of kiss and run events (Lasic et al., 2017). Ryngo 1-23 promotes actin-dependent oligomerization into rings, whereby the authors propose that the bi-directional role of Dyn2 involved a mechanism depending on actin and myosin. We also used Ryngo 1-23 to address how this would influence caveolae, but found no significant effect on caveola dynamics following treatment. This suggests that stimulation of the GTPase activity coupled to actin-dependent oligomerization of Dyn2 was not important for caveola stabilization. Interestingly, when we treated cells with Dyngo 4a, the ratio of Dyn2-positive caveolae was significantly increased suggesting that inhibition of GTP-hydrolysis increases the amount of stable Dyn2 oligomers at caveolae. Furthermore, the treatment resulted in a dramatic increase in caveola stability, supporting the idea that these oligomers stabilized caveolae. However, this impact remained even if dynamin was depleted from the cells. We could show that treatment with this small dynamin effector modulator disrupted the F-actin severely and affected cell morphology in agreement with previous work showing that Dyngo 4a has off-target effects (Park et al., 2013). We propose that Dyngo 4a treatment generally affects PM properties thereby dramatically impacting caveola dynamics, which masks the potential direct effects on Dyn2 activity. A number of studies have used overexpression of Dyn2 mutants such as K44A or treatment with drugs such as Dynasore or Dyngo 4a to restrict caveola budding. These data are consistent with our data, but we propose that this is not due to inhibited GTPase activity but rather the ability of Dyn2 to stabilize caveolae and off-target effects of the drugs.

Caveola fission has been shown to be induced by cholesterol or glycosphingolid accumulation which together with the caveolae coat seem to generate an intrinsically unstable domain prone to fission if not balanced by the restraining force of EHD2 (Hubert et al., 2020). Our data now shows that Dyn2 act independently of EHD2 to also counteract caveolae fission. Most dynamin-like proteins catalyze either membrane fusion or fission processes, but some have also been suggested to work bifunctionally. Similarly, our data suggests that Dyn2 oligomers, known to mediate fission of CCVs, can act to instead restrain caveolae fission and that this does not require GTP hydrolysis. We propose that caveola dynamics are delicately balanced by lipid-driven assembly of the caveola coat and restraining forces produced by regulatory proteins such as EHD2 and Dyn2.

## Materials and Methods

### Reagents

Dyngo 4a (ab120689) and Ryngo 1-23 (ab146050) were both purchased from Abcam. Human Transferrin Alexa Fluor 647-conjugate (T23366) was bought from Invitrogen and SiR-actin (SC0001) from Spirochrome. 1,2-dioleoyl-sn-glycero-3-phosphoethanolamine (DOPE), 1,2-dioleoyl-3-trimethylammonium-propane (chloride salt) (DOTAP), TopFluor-cholesterol (Bodipy-Chol) was purchased from Avanti Polar Lipids Inc (Alabaster, AL, US). DMSO (34943), Chloroform (CHCl_3_), and methanol (MeOH) were purchased from Sigma-Aldrich (St. Louis, MO, US).

### Cell lines and constructs

The HeLa Flp-In T-REx Dynamin2-EGFP-P2A-Caveolin1-mCherry, EGFP-EHD2-P2A-Caveolin1-mCherry and Pacsin2-EGFP-P2A-Caveolin1-mCherry constructs were generated by linearizing pcDNA/FRT/TO/Caveolin1-mCherry (Hubert et al., 2020) with the restriction enzyme HindIII (Thermo Fisher Scientific). The DNA encoding the EGFP-fusion proteins and the P2A peptide was inserted by Gibson assembly using NEBuilder HiFi DNA assembly master mix (New England BioLabs, Ipswich, MA, USA). The HeLa Flp-In T-REx K44A Dynamin2-EGFP-P2A-Caveolin1-mCherry construct was obtained by in vitro mutagenesis of the pcDNA/FRT/TO/Dyn2-EGFP-P2A-Caveolin1-mCherry construct exchanging lysine at position 44 to an alanin using these primers: CCAGAGCGCCGGC<u>GC</u>GAGTTCGGTGCTC and GAGCACCGAACTC<u>GC</u>GCCGGCGCTCTGG. The Flp-In TRex HeLa cell lines were maintained in DMEM supplemented with 10% (v/v) FBS, 100 μg/ml hygromycin B (Thermo Fisher Scientific), and 5 μg/ml blasticidin S HCl (Thermo Fisher Scientific) for plasmid selection at 37°C, 5% CO_2_. Expression at endogenous levels was induced by incubation with 0.5 ng/ml (Cav1-mCh) and 1.0 ng/ml (EGFP-fusion-P2A-Cav1mCh) doxycycline hyclate (Dox, Sigma-Aldrich) for 16–24 h. All cell lines tested negative for mycoplasma. For generation of the ΔNΔEH EHD2-BFP expression vector the ΔNΔEH EHD2 mRNA was subcloned from ΔNΔEH EHD2-mCherry construct (Hoernke et al., 2017) using restriction enzymes XhoI and BamHI (Thermo Fisher Scientific) and inserted into the pTagBFP-C (Evrogen, Moscow, RU) expression vector.

### Fusogenic liposomes

Liposomes were prepared from a lipid mixture of DOPE, DOTAP, and either Bodipy-tagged lipid or unlabeled lipid at a ratio of 47.5:47.5:5. Lipid blends were in MeOH:CHCl_3_ (1:3, v/v). Following the generation of a thin film using a stream of nitrogen gas, the vesicles were formed by addition of 20 mM HEPES (VWR, Stockholm, SE, pH 7.5, final lipid concentration 2.8 μmol/ml) and incubated for 1.5 h at room temperature. Glass beads were added to facilitate rehydration. The liposome dispersion was sonicated for 30 min (Transsonic T 310, Elma, Singen, DE).

### Transfections and cell treatments

Cav1-mCh HeLa cells were transfected with Lipofectamine 2000 (Thermo Fisher Scientific) using Opti-MEM I reduced serum medium (Thermo Fisher Scientific) for transient protein expression. For dynamin2, EHD2 and pacsin2 depletion, Cav1-mCh HeLa cells were transfected with stealth siRNA, specific against human dynamin2, EHD2 /Thermo Fisher Scientific) or ON-TARGETplus siRNA against pacsin2 (Dharmacon) or scrambled control (Thermo Fisher Scientific) using Lipofectamine 2000 and Opti-MEM according to manufacturer’s instructions. Cells were transfected twice over a period of 72 h before the experiment. Protein levels were analyzed by SDS-PAGE and immunoblotting using rabbit anti-Cav1 (ab2910, Abcam), rabbit anti-dynamin2 (PA1-661, Thermo Scientific), rabbit anti-EHD2, RRID:AB_2833022 (Moren et al., 2012), rabbit anti-pacsin2 (2604) and mouse anti-beta-actin (3700) both from Cell Signaling. For drug-treatments cells were treated with 30 μM Dyngo 4a or 1 μM Ryngo 1-23 in live cell medium 30 min respectively 20 min prior to experiment. DMSO (0.001 %) was used as control. For Transferrin uptake, cells were incubated 10 min at 37 °C, 5% CO_2_ with Alexa fluor 647 conjugated Transferrin (5 μg/ml) followed by two washes with ice-cold PBS for a total of 15 min. Cells were immediately fixed with 3% PFA for 10 min at room temperature.

### Fluorescence microscopy analysis

The day prior to imaging cells were induced with Dox and seeded on glass coverslips (CS-25R15) in six-well plates at 3 × 10^5^ cells/well or on precision coverslips (No. 1.5H, Paul Marienfeld GmbH and Co. KG, Lauda-Königshofen, DE) in 24-well plates at 60 × 10^3^ cells/well and incubated at 37 °C, 5% CO_2_. For live-cell microscopy, the media was replaced with live-cell media (DMEM high glucose, no phenol red (Gibco), supplemented with 10% FBS and 1mM sodium pyruvate (Gibco)). For TIRF-microscopy, images were acquired for 5 min at 3 s intervals with each third definitive focus using a Zeiss Axio Observer.Z1 inverted microscope that was equipped with an EMCCD camera iXon Ultra from ANDOR and an alpha Plan-Apochromat TIRF 100×/1.46 oil objective controlled by ZEN software. Confocal stacks were acquired using a Zeiss Cell Observer Spinning Disk Confocal controlled by ZEN interface with an Axio Observer.Z1 inverted microscope, equipped with a CSU-X1A 5000 Spinning Disk Unit and an EMCCD camera iXon Ultra from ANDOR. All micrographs and acquired movies were prepared with Fiji, RRID:SCR_002285 (Schindelin et al., 2012) and Adobe Photoshop CS6, RRID:SCR_014199. Structural illumination microscopy images were acquired using the Zeiss Elyra 7 with Lattice SIM^2^. Images were acquired every 60 to 120 ms as specified by respective figure legend.

### Electron microscopy analysis

3T3-L1 fibroblasts seeded onto glass coverslips were cultured at 37°C in a humidified CO2 incubator. After reaching confluence (approximately 3 days) the 3T3-L1 cells were induced to differentiate and cultured for 10 days as described previously Marsh et al, 1995 PMID 7544796). PM sheets were prepared from the dorsal surface of the differentiated adipocytes by placing the glass coverslips with the cells facing down onto poly-lysine-coated EM grids (Prior et al 2003 PMID 12527752). The grids were then detached from the glass coverslips leaving the dorsal plasma membrane of the cells (PM lawns) attached to the grids. PM lawns were then fixed with 2% paraformaldehyde in PBS at room temperature and washed with PBS. Grids were immunolabelled using primary antibodies to caveolin-1 (BD Biosciences), rabbit anti-EHD2 (Moren et al, 2012) or to dynamin (mouse HUDY1, Sigma), followed by secondary labelling with goat anti-mouse gold and goat anti-rabbit gold. Grids were contrasted using methylcellulose/UA as described previously (Prior et al 2003 PMID 12527752).

### Analysis of caveolae dynamics

Induced HeLa FlpIn cells were imaged on TIRF over 5 min with an acquisition time of 3 s. Imaris software was used for tracking analysis of Cav1-mCh positive structures segmented as spots and structures with a diameter of 0.4 μm as previously described (Mohan et al., 2015). The applied algorithm was based on Brownian motion with max distance travelled of 0.8 μm and a max gap size of 4. The caveolae number per cellular basal membrane was based on spot number identified per frame from the spot segmentation. Colocalization of Dyn2-GFP (wt or K44A) to Cav1-mCh was quantified with Imaris software. Dyn2-GFP positive structures were tracked as described for Cav1-mCh. The first three frames from a TIRF movie was analyzed and a caveolae structure was scored as positive for dynamin if spots from the different channels were present at the same position for at least two frames. A total of 10 cells for each condition was scored and plotted. For caveolae/dynamin segmentation, each Cav1-mCh track within a 5 min TIRF movie from three cells were followed and scored for presences or no presences of Dyn2-GFP. Each track identification number was sorted into two classes; Dyn2-positive or Dyn2-negative and the track duration time and track displacement length was extracted and plotted for each pool. Statistical analysis was performed on track duration (s), track displacement length (μm) and track mean speed (μm/s) data and data are shown as fold change.

### FRAP experiments

Cells were imaged on TIRF and three reference images were recorded before a ROI containing mCh- and GFP-colocalizing structures was photobleached for 1000 ms using maximal laser intensity (488 nm). The fluorescent recovery images were taken every 3 s for a period of 5 min. The intensities of the bleached regions were corrected for background signal and photobleaching of the cell. Data from at least 10 cells were collected per condition and mean FRAP recovery curves were plotted using Prism 5.0 (GraphPad, San Diego, CA, US; RRID:SCR_002798).

### FLIM

For FLIM, HeLa FRT cells were transfected using lipofectamine, or in the case of FlpIn cell lines, induced the day before with 1 nM doxycycline to induce the expression of the protein of interest. Imaging was done on a Leica SP8 FALCON confocal, employing TCSPC (time correlated single photon counting) for the FLIM measurements, which enables high accuracy and high-resolution measurements. Acquisitions were done using a WLL (white light laser) in the GFP excitation area at 40Mhz as a standard and the cooled HyD1 hybrid detector. FRET on relevant pairs was calculated as a reduction of the lifetime in the green fluorophore area. Areas/spots of interest, showing overexpression, was measured using a circular ROI and 10-20 spots were measured per cell and fitted using the LAS X software with the FLIM plugin. Data was plotted using Prism 5.0. FLIM and FRET images were pixel-wise fitted using the LAS X software and exported to tif. Final adjustments were done using Fiji, where images were normalized to each other to show the increase/decrease of lifetime and fret between the images.

### Immunostaining

Induced Dyn2-GFP-Cav1-mCh HeLa cells were transfected with the EHD2-BFP expression vector and seeded on precision coverslips (No. 1.5H, Paul Marienfeld GmbH and Co. KG, Lauda-Königshofen, DE) in 24-well plates at 50 × 10^3^ cells/well and incubated overnight (37°C, 5% CO_2_). Following incubation with Dyngo 4a (30 μM) or DMSO (0.001 %) for 30 min, cells were fixed with 3% PFA in PBS (Electron Microscopy Sciences, Hatfield, PA, US) and subsequent permeabilization and blocking was carried out simultaneously using PBS containing 5% goat serum and 0.05 % saponin. Cells were then immunostained with rabbit anti-PTRF, RRID:AB_88224 (Abcam) followed by goat anti-rabbit IgG secondary antibody coupled to Alexa Fluor 647, RRID:AB_2535814 (Thermo Fisher Scientific) as previously described (Lundmark et al., 2008). Confocal images were acquired using the Zeiss Spinning Disk Confocal microscope (63X lens). Micrographs were prepared using Fiji (Schindelin et al., 2012) and Adobe Photoshop CS6.

### Online supplementary material

Fig. S1 shows the uptake of transferrin-647 in Cav1-mCh HeLa FlpIn cells treated with ctrl or dynamin2 siRNA. Fig. S2 provides additional data on the colocalization of Dyn2-GFP and Cav1-mCh imaged by structured illumination microscopy. Fig. S3 shows Dyn2-GFP localization after EHD2 or pacsin2 depletion and characterization of GFP-EHD2-Cav1-mCh and Pac2-GFP-Cav1-mCh HeLa FlpIn cells. Fig. S4 shows representative FLIM-FRET images of GFP-fusion proteins and Cav1-mCh in FlpIn HeLa cells. Fig. S5 shows the protein titration of K44A Dyn2-GFP and Cav1-mCh cells after Dox-induction in the HeLa FlpIn cell line. Fig. S6 provides additional information of the effect that Dyngo 4a has on the Dyn2-GFP-Cav1-mCh HeLa FlpIn cells. Video 1 shows single particle tracking of Cav1-mCh in a Cav1-mCh HeLa FlpIn cell treated with ctrl siRNA imaged on TIRF. Video 2 shows single particle tracking of Cav1-mCh in a Cav1-mCh HeLa FlpIn cell treated with dynamin2 siRNA imaged on TIRF. Video 3 shows single particle tracking of Cav1-mCh in a Cav1-mCh HeLa FlpIn cell treated with EHD2 siRNA imaged on TIRF. Video 4 shows live cell imaging of a Dyn2-GFP-Cav1-mCh HeLa FlpIn cell using structured illumination microscopy (SIM^2^ algorithm). Video 5 shows live cell TIRF imaging of Dyn2-GFP-Cav1-mCh HeLa FlpIn cells treated 30 min with 30μ of Dyngo 4a.

## Acknowledgements

We acknowledge the Biochemical Imaging Center (BICU) at Umeå University within the National Microscopy Infrastructure, NMI (VR-RFI 2016-00968) and the Microscopy Australia Research Facility at the Center for Microscopy and Microanalysis at the University of Queensland for providing support and assistance with microscopy. We especially thank Irene Martinez at BICU for assistance and expertise with image analysis and data visualization. We thank Claudia Kästner and Abel Pereira da Graça in the ZEISS Democenter in Oberkochen for help with acquiring SIM data. This work was supported by grants from the Swedish Research Council (dnr 2017-04028, dnr 2021-05117 to R.L.) the Swedish Cancer Society (CAN 2017/735, 20 1230 PjF to R.L.), and the National Health and Medical Research Council of Australia (NHMRC) (grant APP1150083 to R.G.P; fellowships APP1156489 to R.G.P.).

## Author contributions

Elin Larsson and Richard Lundmark designed the research and Elin Larsson performed and analysed the molecular biology, live cell imaging and tracking experiments. Kerrie-Ann McMahon and Robert G Parton performed the electron microscopy and Björn Morén performed the FLIM experiments. Elin Larsson and Richard Lundmark conceived the work and wrote the manuscript and all authors contributed with text, figures and editing.

