## Supplementary material for "Dynamin2 stabilizes plasma membrane-connected caveolae by restraining fission"

**Affiliations**
1. Integrative Medical Biology, Umeå University, 901 87, Umeå, Sweden.
2. Institute for Molecular Bioscience, The University of Queensland, Brisbane, QLD, Australia
3. Centre for Microscopy and Microanalysis, The University of Queensland, Brisbane, QLD, Australia

**This PDF file includes:**

Figs. S1 to S6 and legends to Movie 1 – 5.

Supplemental Fig. Legends

Figure S1

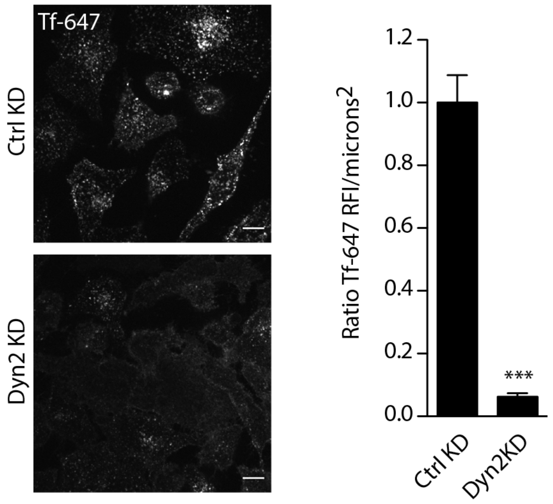


**Fig. S1. Transferrin uptake in control and Dynamin2 depleted HeLa Cav1-mCh FlpIn cells.** Representative images of fixed Cav1-mCh cells transfected with siRNA as indicated for 72 hours before 10 min incubation with Alexa fluor 647 conjugated Transferrin (5 µg/ml) at 37°C. Quantification show relative Tf-647 fluorescent intensity/microns$^2$. Significance was assessed using *t* test, *** p ≤0.001.

Figure S2

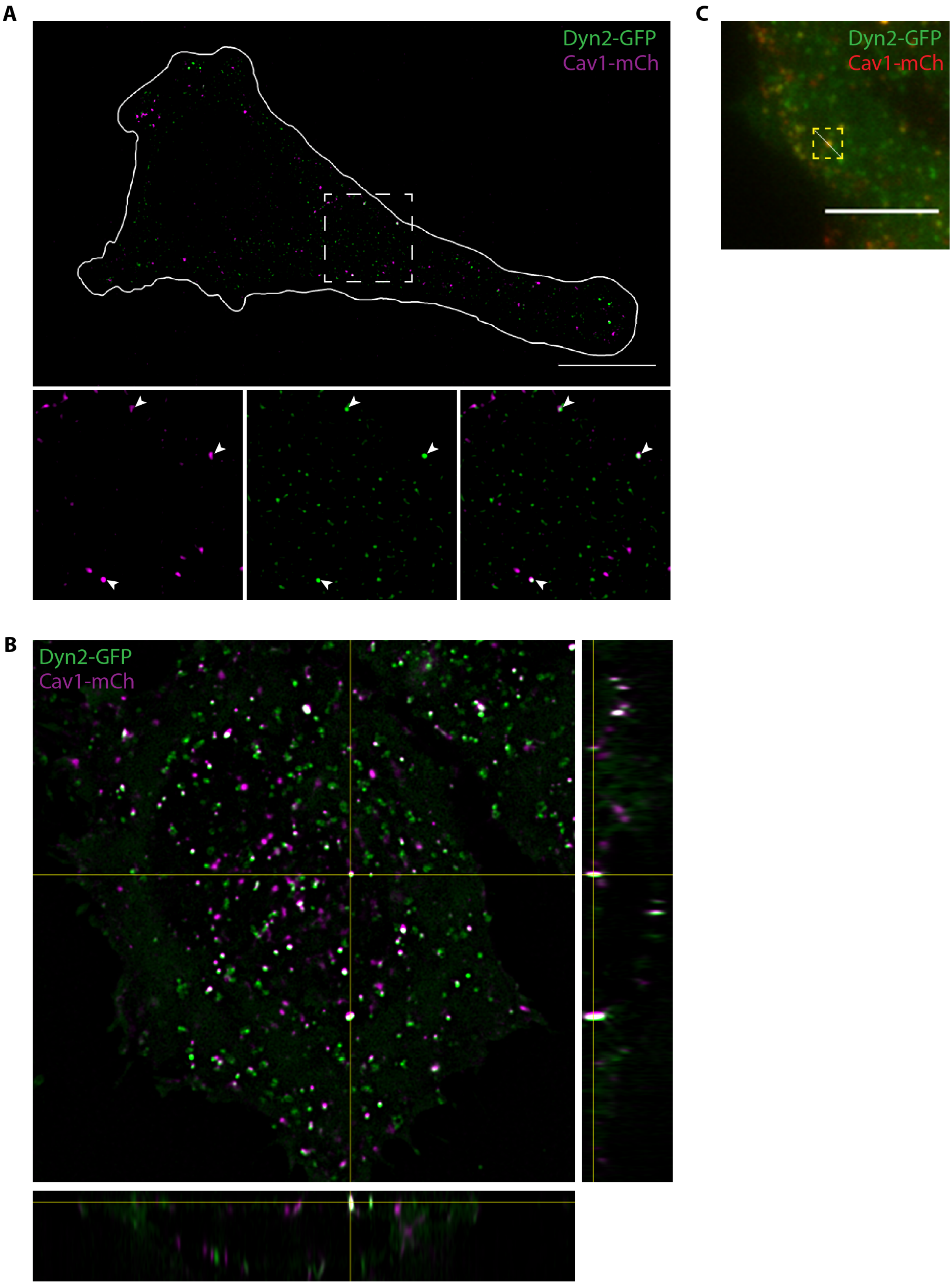


**Fig. S2. Dyn2-GFP colocalizes with Cav1-mCh. (A)** Representative SIM$^2$ image of Dyn2-GFP-Cav1-mCh cell imaged live. White line marks the outline of the cell and dashed square marks location of the magnified areas in the bottom panel. White arrows highlight structures positive for both Dyn2-GFP and Cav1-mCh. Scale bar, 10 µm. **(B)** Orthogonal view of a live Dyn2-GFP-Cav1-mCh cell imaged with Lattice SIM[2]. **(C)** Dyn2-GFP-Cav1-mCh cell with line which indicate location of kymograph in Fig. 2D. Scale bar, 5 µm.

Figure S3

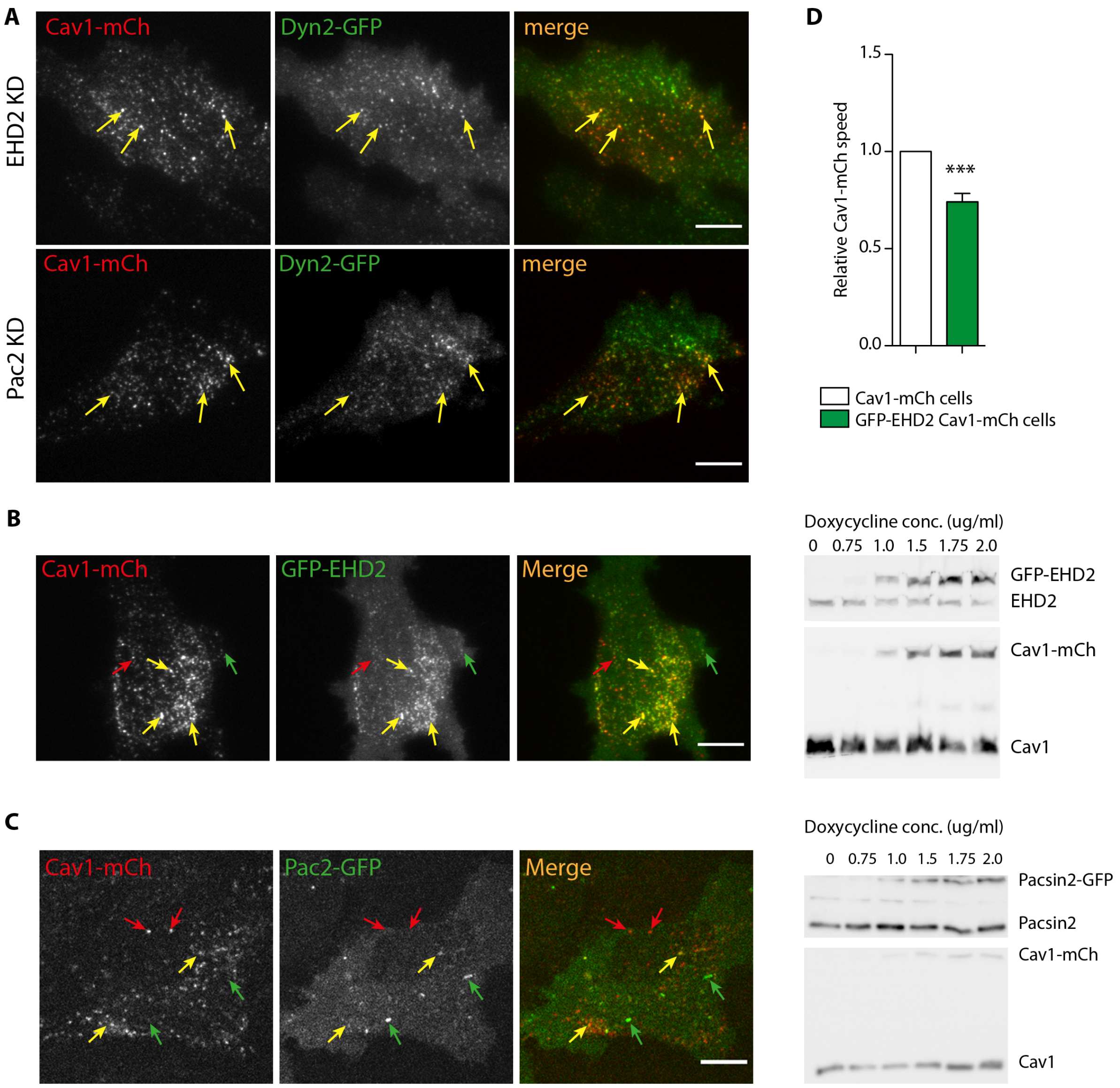


**Fig. S3. Dyn2-GFP localization after EHD2 or Pac2 depletion and HeLa FlpIn characterization of GFP-EHD2-Cav1-mCh and Pac2-GFP-Cav1-mCh cells. (A)** Representative image from TIRF movie of Dyn2-GFP-Cav1-mCh cells treated with siRNA directed against EHD2 or pacsin2 as indicated. Yellow arrows highlight caveolae positive for Dyn2-GFP. **(B)** Representative image from TIRF movie of a GFP-EHD2-Cav1-mCh cell. Red arrow depicts structures only positive for Cav1-mCh, green arrow highlights spots only positive for GFP-EHD2 and yellow arrows highlight structures positive for both GFP-EHD2 and Cav1-mCh. Immunoblot of GFP-EHD2-Cav1-mCh cells treated with concentrations of doxycycline as indicated. Doxycycline concentration used for further experiments were 1.0 ng/ml. **(C)** Representative image from TIRF movie of Pac2-GFP-Cav1-mCh cell. Red arrow depicts structures only positive for Cav1-mCh, green arrow highlights spots only positive for Pac2-GFP and yellow arrows highlight structure positive for both Pac2-GFP and Cav1-mCh. Immunoblot of Pac2-GFP-Cav1-mCh cells treated with concentrations of doxycycline as indicated. Doxycycline concentration used for further experiments were 1.0 ng/ml **(D)** Quantification of Cav1-mCh track mean speed in GFP-EHD2-Cav1-mCh cells (green). Numbers were related to Cav1-mCh cells (white). Track mean ± SEM from at least 7 cells per condition are shown. Significance was assessed using *t* test, *** p ≤0.001. All scale bars, 10 µm.

Figure S4

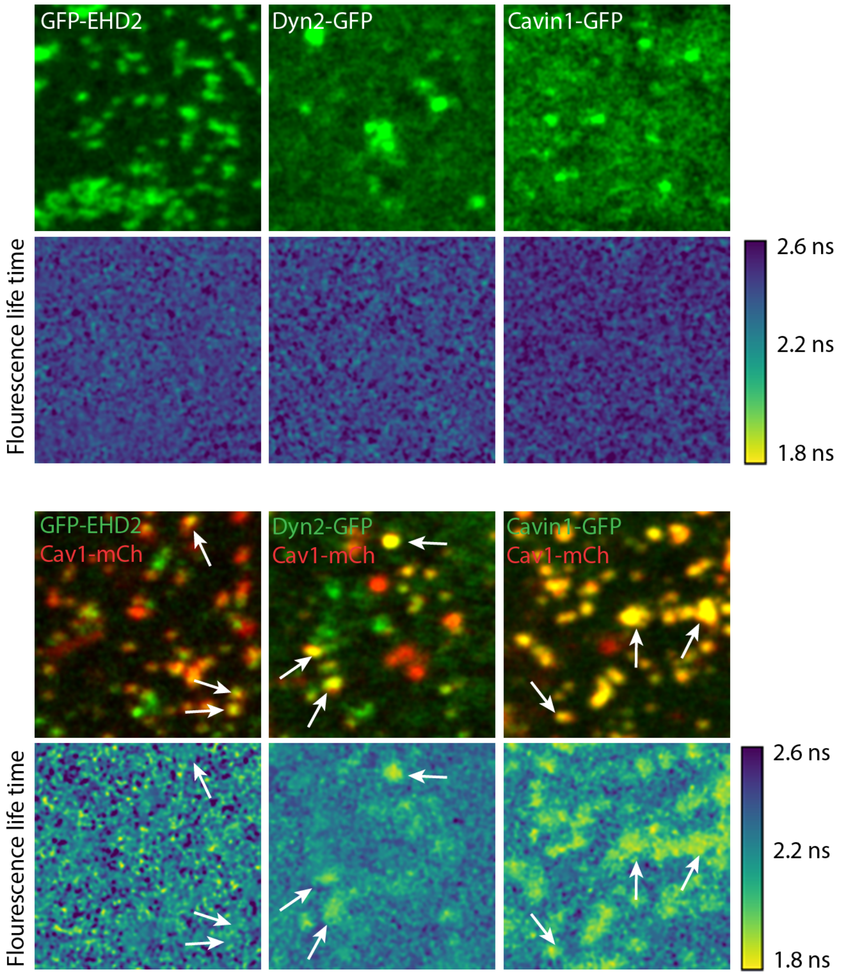

**Fig. S4. FLIM-FRET of GFP fusion proteins.** Top of the two panels show representative images of cells expressing fusion proteins as indicated. Bottom of the two panels show EGFP fluorescence lifetime. White arrows highlight structures where EGFP and mCherry colocalize.

Figure S5

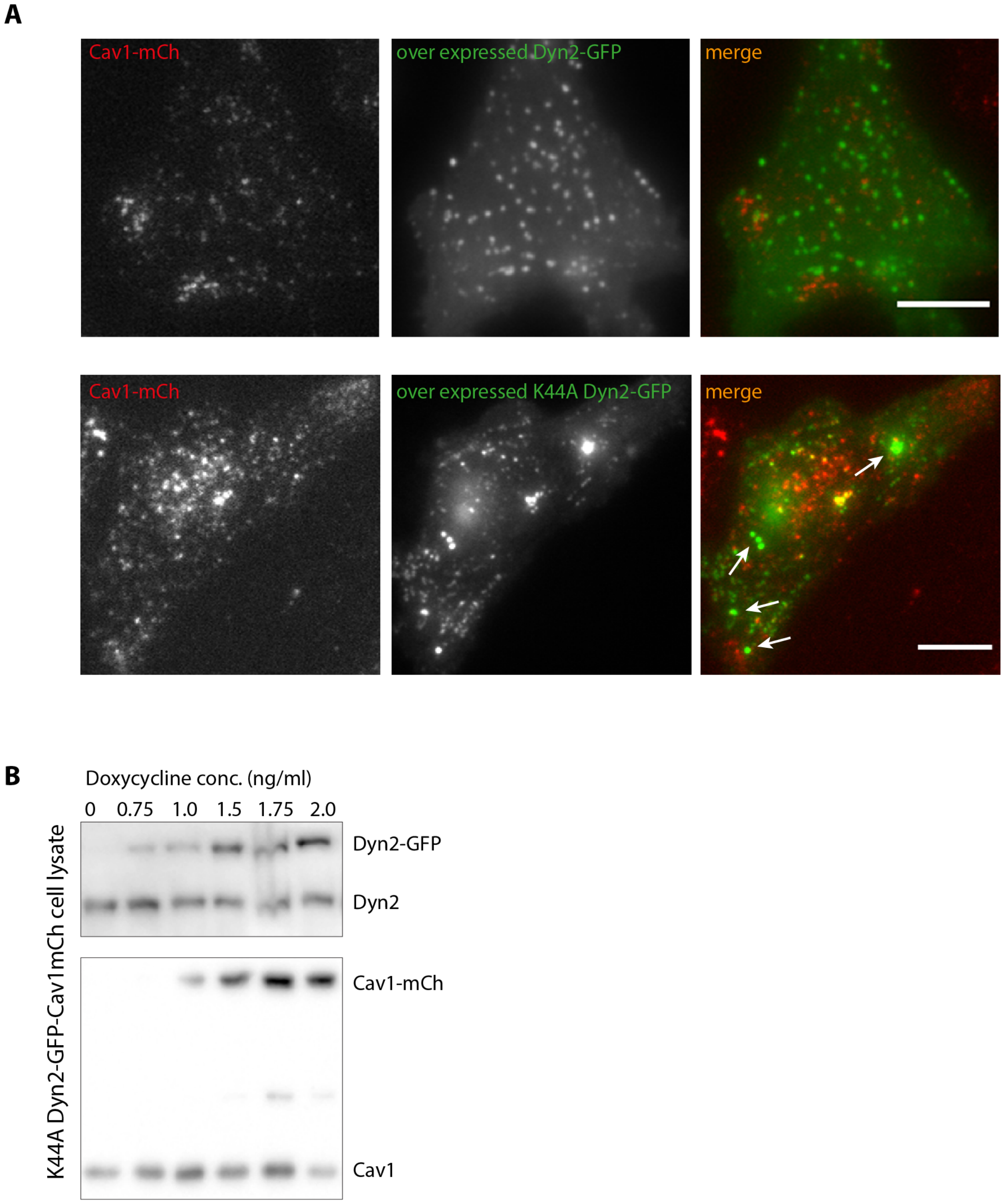


**Fig. S5. Over expression of dynamin2 cause abnormal protein aggregation in Cav1-mCh HeLa FlpIn cells. (A)** Representative images of Cav1-mCh HeLa FlpIn cells transiently expressing wild type Dyn2-GFP (top panel) or K44A Dyn2-GFP (bottom panel). **(B)** Immunoblot of K44A Dyn2-GFP-Cav1-mCh cells treated with concentrations of doxycycline as indicated. Doxycycline concentration used for experiments were 1.0 ng/ml.

Figure S6

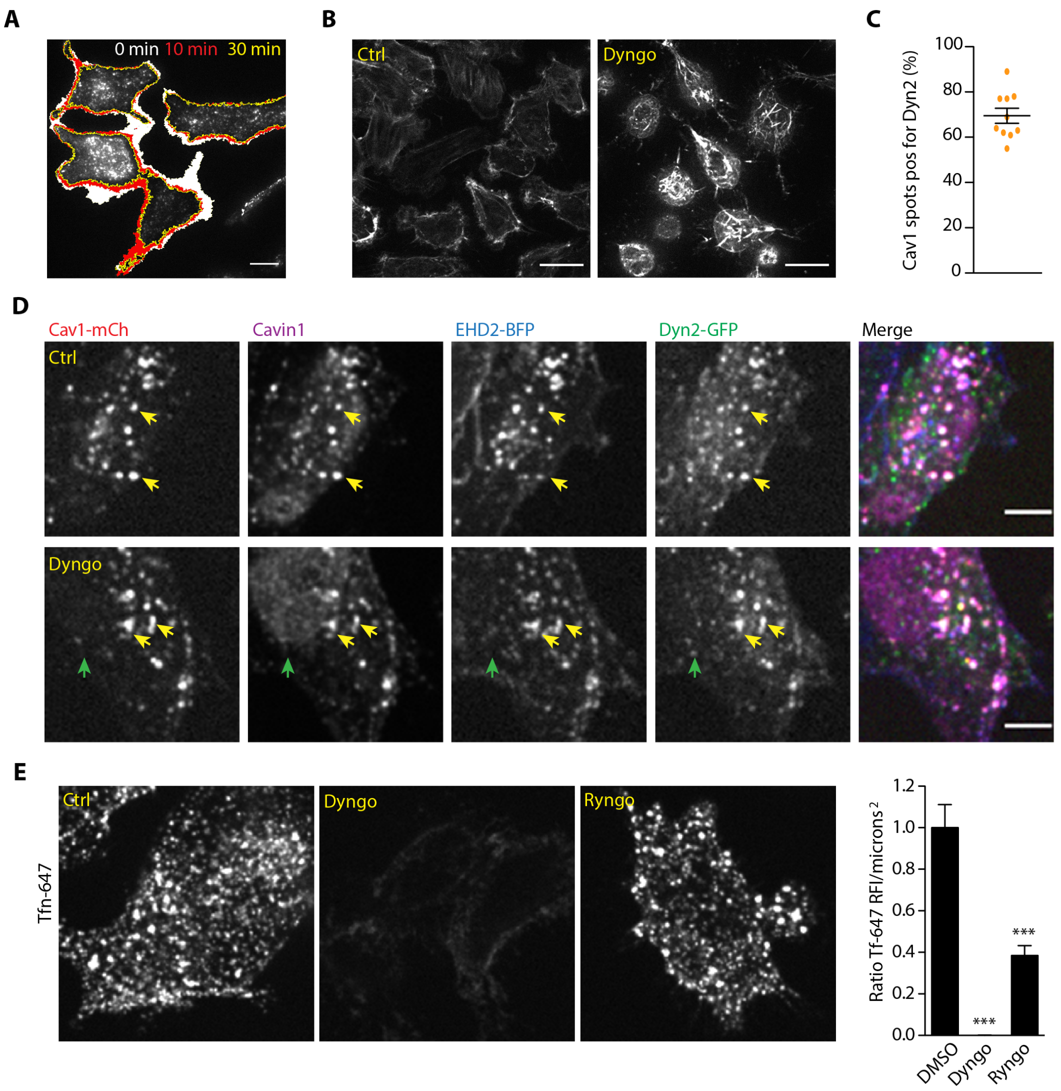


**Fig. S6. Dyngo 4a treatment affects cell morphology and F-actin. (A)** Example of basal plasma membrane retraction after Dyngo 4a addition. White area depicts the area retracted between time point 0 min to 10 min. Red area depicts the area retracted between time point 10 min to 30 min. Yellow outline illustrates the basal plasma membrane surface area after 30 min treatment with Dyngo 4a. Scale bar, 10 µm. **(B)** Representative confocal image of basal membrane in Dyn2-GFP-Cav1-mCh cells treated with DMSO (Ctrl) or Dyngo 4a for 30 minutes and incubated with the F-actin marker SiR-actin. Scale bar, 20 µm. **(C)** Quantification of percentage of caveolae positive for Dyn2-GFP after 30 min Dyngo 4a treatment. Data are shown as scatter dot plot, mean ± SEM. **(D)** Representative immunofluorescent staining of

Dyn2-GFP-Cav1-mCh cells. Cells transiently expressing EHD2-BFP were treated with DMSO (Ctrl) or Dyngo 4a for 30 minutes and then fixed and stained with cavin1-antibody (magenta). Yellow arrows depict colocalizing structures and green arrow depicts only Dyn2-GFP positive structure. Scale bar, 10 μm. **(E)** Representative image of fixed Dyn2-GFP-Cav1-mCh cells treated with DMSO (Ctrl), Dyngo 4a or Ryngo before 10 min incubation with Alexa fluor 647 conjugated Transferrin (5 μg/ml) at 37 °C. Quantification show relative Tf-647 fluorescent intensity/microns$^2$. Significance was assessed using *t* test, *** p ≤0.001.

**Video 1. Single particle tracking of Cav1-mCh in a HeLa Cav1-mCh FlpIn cell.** Representative single particle tracking of Cav1-mCh structures of a HeLa Cav1-mCh cell treated with ctrl siRNA imaged on TIRF every 3$^{rd}$ second for 5 min. The mCh fluorescence was segmented using Imaris 9.5.1 and white spheres represents caveolae positive for Cav1-mCh. Rainbow-colored trajectories display duration time and displacement length of the caveolae as they are being traced. Purple trajectories represent short duration times (9 - 15 s) whereas red trajectories represent caveolae with long duration times (291 -300 s).

**Video 2. Dynamin2 depletion increases caveola dynamics.** Representative single particle tracking of Cav1-mCh structures of a HeLa Cav1-mCh cell treated with dynamin2 siRNA imaged on TIRF every 3$^{rd}$ second for 5 min. The mCh fluorescence was segmented using Imaris 9.5.1 and white spheres represents caveolae positive for Cav1-mCh. Rainbow-colored trajectories display duration time and displacement length of the caveolae as they are being traced. Purple trajectories represent short duration times (9 - 15 s) whereas red trajectories represent caveolae with long duration times (291 -300 s).

**Video 3. EHD2 depletion increases caveola dynamics.** Representative single particle tracking of Cav1-mCh structures of a HeLa Cav1-mCh cell treated with EHD2 siRNA imaged on TIRF every 3$^{rd}$ second for 5 min. The mCh fluorescence was segmented using Imaris 9.5.1 and white spheres represents caveolae positive for Cav1-mCh. Rainbow-colored trajectories display duration time and displacement length of the caveolae as they are being traced. Purple trajectories represent short duration times (9 - 15 s) whereas red trajectories represent caveolae with long duration times (291 -300 s).

**Video 4. Localization of Dyn2-GFP to caveolae.** Dyn2-GFP-Cav1-mCh cells were induced with 1ng/ml Dox and imaged using structured illumination microscopy (SIM$^2$ algorithm). Cells were imaged every 60 ms for a total of 100 frames. Green channel represents Dyn2-GFP and magenta represents Cav1-mCh.

**Video 5. Dyngo 4a treatment results in static caveolae within the plasma membrane.** Representative live cell movie of Dyn2-GFP-Cav1-mCh FlpIn cells treated with Dyngo 4a for 30 min prior to imaging. Cells were imaged on TIRF every 3$^{rd}$ second for 5 min. Cav1-mCh is seen in red and Dyn2-GFP in green.
